# Dichotomous SMAD2/3 regulation and selective anti-hypertrophic activity of heparin during in vitro chondrogenesis of mesenchymal stromal cells

**DOI:** 10.1101/2025.09.02.673657

**Authors:** Sven Schmidt, Safak Chasan, Helen F. Dietmar, Felicia A.M. Klampfleuthner, Tilman Walker, Uwe Freudenberg, Wiltrud Richter, Solvig Diederichs

**Author notes:** Corresponding author: Address for correspondence PD Dr. Solvig Diederichs, Research Centre for Molecular and Regenerative Orthopaedics Department for Orthopaedics, Medical Faculty of the Heidelberg University Schlierbacher Landstraße 200a, 69118 Heidelberg, Germany.

## Abstract

**Background:** Endochondral instead of chondral differentiation hinders mesenchymal stromal cell (MSC) application for clinical cartilage regeneration. We previously showed that heparin-polyethylene glycol (PEG) hydrogels loaded with transforming growth factor beta (TGF-β) instructed stable chondral MSC development in vivo. We here assessed this approach in vitro, utilizing heparin-PEG hydrogels or soluble heparin supplementation of chondrogenic medium.

**Methods:** Human MSCs were cultured in heparin-PEG hydrogels (22.4 mg/mL crosslinked heparin, 120ng TGF-β1) or as hydrogel-free pellet cultures treated with soluble heparin (0, 10, 100, 700 μg/mL) in TGF-β1-containing (10 ng/mL) chondrogenic medium. Chondral and endochondral signaling (1–3 h, 4 weeks) and cartilage matrix formation (4 weeks) were analyzed using Western blot, histology, qPCR, ELISA, and enzyme activity.

**Results:** Unlike in vivo, human MSCs differentiated in heparin-PEG hydrogels into type X collagen and alkaline phosphatase-positive hypertrophic chondrocytes in vitro. Interestingly, treatment with soluble heparin (10-700 µg/mL) revealed reduced TGF-β-small mother against decapentaplegic (SMAD)3 but not SMAD2 activation at unaffected type II collagen and proteoglycan/DNA levels. We propose that the stimulation of the insulin-AKT pathway by heparin aided in maintaining SMAD2 activation which apparently plays a more prominent role than SMAD3 for MSC chondrogenesis. Heparin treatment inhibited the pro-hypertrophic WNT/β-catenin pathway in vitro but insufficiently silenced TGF-β-SMAD1/5/9 activation and unfortunately reduced anti-hypertrophic prostaglandin E2 (PGE2) levels. Ultimately, treatment with 10 µg/mL heparin reduced the upregulation of several hypertrophy markers (*MEF2C, IHH, IBSP* mRNAs, alkaline phosphatase activity) below control levels, but type X collagen remained unresponsive. Thus, soluble heparin treatment was similarly selective and effective as previous anti-hypertrophic interventions (parathyroid-hormone related protein (PTHrP)-pulses, wingless-int (WNT)-inhibition), while offering technical simplicity, reduced cost, and solvent-free formulation.

**Conclusions:** Taken together, heparin-TGF-β showed a novel dichotomous SMAD2/3 inhibition at maintained chondrogenic power and context-dependent lineage-instructive properties: permitting endochondral differentiation in vitro but chondral development in vivo. Thus, environmental contributions are mandatory to allow heparin-PEG-guided chondral versus endochondral lineage commitment of MSCs in vivo, potentially involving SMAD1/5/9 suppressors and PGE2 sources.

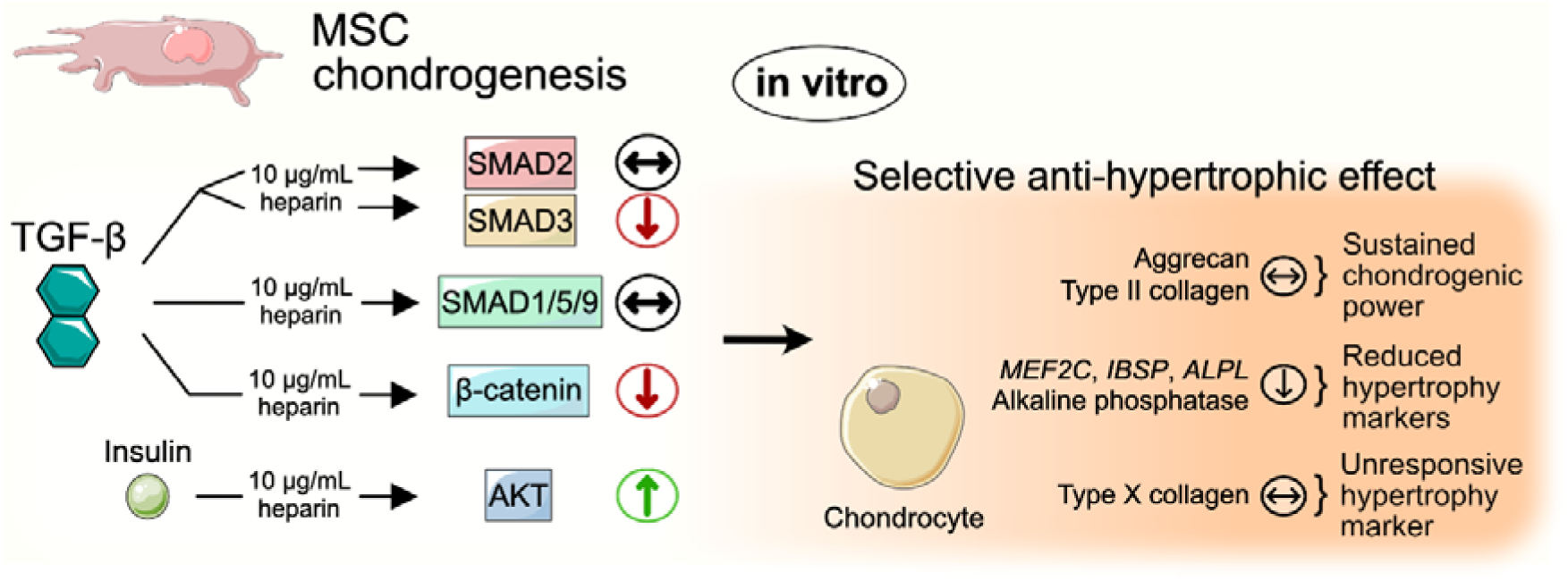
Graphical Abstract.

## Introduction

Mesenchymal stromal cells (MSCs) are a promising autologous chondrocyte source for regenerating cartilage tissues. Cartilage regenerative strategies often involve in vitro chondrogenic MSC pre-differentiation to avoid dependence on uncontrollable local in vivo signals for the directed differentiation into chondrocytes and the in-situ formation of neocartilage. Application of pre-differentiated rather than non-differentiated MSC-based cartilage grafts can significantly enhance extracellular matrix deposition and improve repair quality [1, 2].

Standard MSC in vitro chondrogenesis conditions employ the essential chondro-inductive transforming growth factor TGF-β which activates the pro-chondrogenic small mother against decapentaplegic (SMAD)2 and SMAD3 [3]; and insulin/AKT signaling cooperates in activating SMAD2 [4]. SMAD2/3 induce the chondrogenic master transcription factor SRY-box transcription factor (SOX)9, initiating the secretion of the hallmark extracellular cartilage matrix proteins type II collagen and the proteoglycan aggrecan [3].

Yet under these conditions, MSCs unavoidably commit to the endochondral instead of the desired chondral cell lineage in vitro and phenocopy the endochondral bone development and growth of the skeleton [3]. Cells develop into hypertrophic chondrocytes that additionally release proteins associated with matrix mineralization and bone formation, such as type X collagen, alkaline phosphatase (ALP), and integrin-binding sialoprotein (IBSP) [5, 6, 7]. In vivo, the resulting hypertrophic neocartilage is prone to mineralize and to remodel into bone tissue [5], a highly undesired outcome for cartilage regeneration.

The current understanding of the signals driving the hypertrophic MSC misdifferentiation involves a complicated interplay between TGF-β-SMAD1/5/9 activity, cell-autonomous wingless-int (WNT) signaling, and dysregulation of the feedback loop between parathyroid-hormone related protein (PTHrP) and Indian Hedgehog (IHH) [8, 9, 10]. At the initiation of chondrogenesis, TGF-β non-canonically activates SMAD1/5/9 [11], that is associated with bone morphogenetic protein signaling which promotes chondrocyte hypertrophy in the growth plate [12]. Additionally, TGF-β stimulates β-catenin accumulation in MSCs undergoing chondrogenesis [13], the mediator of canonical WNT signaling. Knockout and inhibitor studies demonstrated that WNT activity drives chondrocyte hypertrophy in both the growth plate and MSC chondrogenesis [8, 14, 15]. Furthermore, once PTHrP expression is downregulated during advancing MSC chondrogenesis, WNT signaling contributes to IHH upregulation [8], a prominent driver of endochondral ossification in the growth plate [16].

In line with these insights, WNT inhibition and pulsed PTHrP treatment are currently the most effective anti-hypertrophic interventions during MSC chondrogenesis [8, 15, 17]. Along with a strong reduction of ALP protein levels by 80% and more, WNT inhibition and PTHrP pulses reduce the expression of many hypertrophy markers including *IBSP* and *IHH* messenger ribonucleic acids (mRNAs). However, type X collagen deposition remained unaffected, and the generated neocartilage mineralized and was remodeled into bone tissue after subcutaneous implantation, indicating a still insufficient instruction and stabilization of MSC commitment into the chondral cell lineage.

A major success in directing MSCs into the desired chondral lineage was to harness the lineage-determining capacity of sulfated glycosaminoglycans (sGAGs) of the heparan sulfate family on stem and progenitor cell development [18]. The unique structural features of heparan sulfates, most importantly their sulfation, enable them to interact with growth factors, morphogens, and cell surface receptors. Heparin is the most highly charged heparan sulfate and has a strong affinity to TGF-β [19, 20]. By utilizing heparin to immobilize TGF-β in a novel biodegradable poly(ethylene glycol) (PEG)-based hydrogel, we could instruct MSC commitment in vivo into the chondral lineage for the first time [18]. Allowing long-term retention of TGF-β and silencing WNT/β-catenin and SMAD1/5/9 pathways, heparin facilitated the formation of permanent neocartilage with high levels of type II collagen, while remaining remarkably free of type X collagen and long-term resistant to calcification. However, this was achieved in the undefined environment of a subcutaneous mouse model, and it remained unclear whether a similar success can be replicated under the defined conditions of in vitro chondrogenesis.

In our previous in vitro experiments with heparin-PEG hydrogels, MSC chondrogenesis resulted in mostly pericellular cartilage matrix deposition, significantly lower than that of articular chondrocytes [21]. Excessive TGF-β immobilization in these hydrogels likely limited chondrogenic MSC differentiation, thus confounding the analysis of chondrocyte hypertrophy. By contrast, robust chondrogenic differentiation and homogeneous matrix deposition was achieved from goat MSCs or human fetal chondroprogenitor cells when supplying TGF-β exogenously to heparin-PEG-gelatin or heparin-hyaluronan hydrogels, respectively [22, 23]. Levinson et al. reported positive type X collagen staining in their generated neocartilage, but a potential regulation by heparin remained unclear owing to a lack of heparin-free controls [23]. Importantly, our hydrogels incorporated over 20-fold more crosslinked heparin (2.24% vs.0.1% w/v) and may thus be more effective in suppressing type X collagen production.

Heparin has alternatively been supplied to chondrogenic cultures as soluble salt. Early studies using chick limb bud mesenchymal cells reported that low heparin concentrations (1-10 μg/mL) can stimulate proteoglycan synthesis, while higher concentrations (200 μg/mL) were detrimental [24, 25]. While more recent studies confirmed the compatibility of adequate soluble heparin doses with chondrogenesis of bone marrow or adipose-derived MSCs [26, 27, 28], effects on chondrocyte hypertrophy and specifically type X collagen deposition have not been addressed. Interestingly, soluble heparin reduced growth and differentiation factor (GDF)5-induced ALP activity in chondrogenic ATDC5 cultures [29], supporting a potential anti-hypertrophic effect of heparin in vitro.

The aim of this study was to investigate whether heparin-PEG hydrogels or heparin in soluble form can support cell fate instruction during TGF-β-driven MSC in vitro chondrogenesis into the chondral lineage to form stable (non-hypertrophic) neocartilage tissues. To this end, we immobilized MSCs in the previously established TGF-β1-loaded heparin-PEG hydrogels [18] and cultured these in chondrogenic medium containing additional free TGF-β1. Moreover, hydrogel-free MSC pellets were cultured in standard chondrogenic medium supplemented with soluble heparin and TGF-β1. We investigated the neocartilage formed after four weeks for hallmark characteristics of chondrocyte hypertrophy including type X collagen deposition, and investigated the influence of soluble heparin treatment on the activation of pro-chondrogenic TGF-β-SMAD2/3 and insulin-AKT pathways, as well as of pro-hypertrophic WNT/β-catenin and SMAD1/5/9 pathways.

## Materials and Methods

### Isolation, expansion, and chondrogenic differentiation of human MSCs from bone marrow

Adhering to the ethical guidelines as detailed below and after written informed consent of patients (28-83 years) undergoing total hip endoprosthesis surgery, human MSCs were isolated from bone marrow samples. A Ficoll-Paque™ PLUS (Cytiva, Freiburg, Germany) density gradient was used for mononuclear cell enrichment. Cells were then cultivated in 0.1% gelatin-coated culture flasks (37°C, 6% CO_2_) using an expansion medium composed of high glucose (4.5 g/L) Dulbecco’s Modified Eagle’s Medium (DMEM) supplemented with 12.5% fetal bovine serum, 1% penicillin/streptomycin (Pen Strep, Gibco™, Thermo Fisher Scientific, Darmstadt, Germany), 2 mM L-glutamine, 1% non-essential amino acids (Minimum Essential Medium), 1% 2-mercaptoethanol (all Gibco™, Thermo Fisher), and 4 ng/mL fibroblast growth factor-2 (Active Bioscience, Hamburg, Germany; Miltenyi Biotec, Bergisch Gladbach, Deutschland). After 24 hours, nonadherent cells were removed by washing with phosphate-buffered saline (PBS), and the medium was exchanged thrice weekly.

Confluent passage 3 cells were subjected to pellet culture (5×10^5^ MSCs) in chondrogenic medium containing high glucose DMEM, 1% Pen Strep, 0.1 μM dexamethasone, 0.17 mM L-ascorbic acid-2-phosphate, 2 mM sodium pyruvate, 0.35 mM proline (all Sigma-Aldrich, Darmstadt, Germany), 1% ITS™+ Premix (Corning Life Sciences, New York City, USA), and 10 ng/mL TGF-β1 (PeproTech, Darmstadt, Germany), supplemented as indicated with heparin (10, 100, 700 μg/mL; sodium salt from porcine mucosa; MW: ≈15,000 Da; Calbiochem, Merck Millipore, Darmstadt, Germany) or chondroitin-4-sulfate (10, 100, 700 μg/mL; sodium salt from bovine trachea; Sigma-Aldrich). Medium exchanges were performed three times per week.

### Preparation of heparin-PEG hydrogels

The heparin-PEG hydrogels were produced as previously established [21, 30]. To achieve a hydrogel with 50% matrix metallopeptidase (MMP)-sensitive linkers, 0.75 mol non-MMP-sensitive thiol end-functionalized starPEG, 0.75 mol thiol end-functionalized starPEG-MMP conjugates, and 1 mol maleimide-functionalized heparin were reconstituted in PBS and mixed with 1.2 × 10^6^ passage 3 MSCs and TGF-β1 (120 ng per 60 µL hydrogel construct). The hydrogels were then polymerized in 6 mm × 2 mm disc moulds and cultured in chondrogenic medium for 28 days. Since the hydrogels swell in physiological salt solutions, the prepared 22.4 mg/mL heparin in the hydrogel formulation (1.344 mg per 60 µL hydrogel) resulted in ∼18 mg/mL heparin in cultured gels. Medium was exchanged thrice weekly.

### Protein extraction

For collagen extraction, one day 28 pellet per treatment group and donor was digested for 16 h using pepsin (2.5 mg/mL pepsin, 0.5 M acetic acid, 0.2 M NaCl, Carl Roth, Karlsruhe, Germany). After adjusting the pH to 7 using 1 M Tris base (Carl Roth), the collagens were extracted with 4.5 M NaCl (overnight, 4°C). Of note, this procedure also degrades all typical reference proteins, including β-actin. Collagens were precipitated for 4 h at –20 °C using 100% ethanol. Pelleted collagens were then resuspended in lysis buffer (50mM Tris, 150mM NaCl, 1% Triton™ X-100, Sigma-Aldrich).

Whole cell lysates were prepared from 1-2 pellets per treatment group and donor using PhosphoSafe™ Extraction Reagent (Merck Millipore) supplemented with 1 mM Pefablock^®^ SC (Sigma-Aldrich) for the detection of SMAD and AKT proteins. For the detection of cytosolic β-catenin, cells were treated with Saponin buffer (0.05% in Tris-buffered saline and 1 mM MgCl_2_, both Sigma-Aldrich) supplemented with 1 mM Halt™ Proteinase/Phosphatase Inhibitor Cocktail (Thermo Fisher Scientific). The extracted supernatant was mixed with Laemmli buffer (33.2% (w/v) glycerol, 249 mM Tris-HCl pH 6.8 (both Carl Roth), 8.0% (w/v) sodium dodecyl sulfate (SDS), and 0.02% bromophenol blue (both Sigma-Aldrich). Samples were boiled for 5 min at 95°C under constant agitation.

### SDS-PAGE and Western blotting

Collagens were separated using 6% polyacrylamide gels, and all other proteins using 10% gels following standard gel electrophoresis protocols. The proteins were then transferred onto a nitrocellulose membrane (Amersham™, GE Healthcare, Chalfont St Giles, United Kingdom). Membranes were cut horizontally to detect proteins of different sizes. For collagen detection, the lower part of the membrane was incubated with mouse anti-human type X collagen antibody (1:500, clone X53, #41-9771-82, Invitrogen) and the upper part with mouse anti-human type II collagen antibody (1:1000, clone II-4C11, #63171, MP Biomedicals, Eschwege, Germany). For the detection of SMAD, AKT, or β-catenin, the upper membrane sections were treated with rat monoclonal anti-pSMAD1/9 (1:1000, pS463/pS465, pS465/pS467, clone N6-1233 (RUO), #562509, BD Biosciences, East Rutherford, NJ, USA), rabbit monoclonal anti-pSMAD2 (1:500, pS465/pS467, clone 138D4, #3108), rabbit monoclonal anti-pSMAD3 (1:500, pS423/pS425, clone C25A9, #9520), rabbit polyclonal anti-pAKT (1:500, pS473, #9721, all Cell Signaling Technology, Danvers, MA, USA) or mouse monoclonal anti-β-catenin antibodies (1:10,000, clone 14/Beta-Catenin (RUO), #610154, BD Biosciences). The lower membrane sections were probed using mouse monoclonal anti-β-actin (1:10,000, clone AC-15, #GTX26276, GeneTex, Irvine, CA, USA), or mouse monoclonal anti-CD81 (1:500, clone B-11, #sc-166029, Santa Cruz Biotechnology, Dallas, TX, USA). For total SMAD detection, membranes were re-incubated with rabbit monoclonal anti-SMAD1/5 (SMAD1: 1:500, clone EP565Y, #ab33902; SMAD5: 1:1000, EP619Y, #ab40771; both Abcam, Berlin, Germany) or rabbit monoclonal anti-SMAD2/3 (1:250, clone D7G7, #8685), and for total AKT detection with rabbit polyclonal anti-AKT (1:100, #9272, both Cell Signaling Technology). Secondary antibodies were HRP-conjugated goat anti-rat antibody (1:1000, HAF005, Bio-Techne, R&D Systems, Minneapolis, MN, USA), peroxidase-coupled goat anti-mouse antibody (1:10000, #115-035-071), or peroxidase-coupled goat anti-rabbit antibody (1:10000, #111-035-046, both Jackson ImmunoResearch Laboratories, West Grove, PA, USA). All primary and secondary antibodies were diluted in 5% skim milk in TBS-T (primary: incubated overnight at 4°C, secondary: 2 h at room temperature). EnhancedChemiLuminescence (ECL) solutions (Roche, Mannheim, Germany) were used for band visualization.

### Type II collagen enzyme-linked immunosorbent assay (ELISA)

The Type II Collagen Detection kit (Chondrex, Woodinville, WA, USA) was used to quantify the type II collagen content in the collagen extraction samples, following the manufacturer’s instructions.

### Glycosaminoglycan and DNA quantification

Per treatment group and donor, 1 or 2 day 28 pellets were digested with 0.1 mg/mL proteinase K (in Tris-HCl (50 mM, 1 mM CaCl_2_, Sigma-Aldrich), Thermo Fisher) for 18 h at 60°C with continuous shaking. Following dilution in Tris/EDTA buffer (Invitrogen, Thermo Fisher Scientific), samples were mixed with 1,9-dimethyl methylene blue solution (48 μM, Thermo Fisher Scientific, in 40 mM NaCl, 40 mM glycine, both Carl Roth), and the absorbance was measured at 530 nm. The values obtained were referred to a chondroitin 6-sulfate standard curve.

The Quant-iT™-PicoGreen® kit (Invitrogen) was used according to the manufacturer’s instructions to quantify the DNA content in the proteinase K-digested samples. The resulting fluorescence values were compared to a λ-DNA-derived standard curve.

### Prostaglandin E2 ELISA

For quantification of the PGE2 content, 48-hour conditioned culture supernatants from 7-9 pellets per treatment group and donor were pooled at the indicated time points. Following the provider’s protocol, a colorimetric competitive PGE2 ELISA (Enzo Life Sciences, Farmingdale, NY, USA) was performed.

### ALP enzyme activity assay

The supernatants from 7-9 pellets per treatment group and donor were pooled at the indicated time points. An equal volume of p-nitrophenyl phosphate substrate (10 mg/mL in 0.1 M glycine, Carl Roth, 1 mM MgCl_2_, and 1 mM ZnCl_2_, Sigma-Aldrich, pH 9.6) was added. After 120 minutes the substrate conversion was detected at 405 nm and corrected by the signal at 490 nm. The enzyme activity was calculated using a standard curve derived from p-nitrophenol (Sigma-Aldrich).

### Analysis of gene expression

Three day 28 pellets were pooled and homogenized for each group and donor using a polytron. RNA was extracted from the homogenate using peqGOLD TriFast™ reagent (Peqlab, VWR, Avantor, Erlangen, Germany) following the manufacturer’s instructions. Complementary DNA was synthesized from 500 ng total RNA using oligo-dT primers and the OmniScript^®^ Reverse Transcription Kit (QIAGEN, Hilden, Germany). SYBR^®^ Green master mix (Thermo Fisher), specific primer sequences (Supplementary Table S1), and a LightCycler^®^ 96 device (Roche Diagnostics, Basel, Switzerland) were used to determine the transcript levels of the genes of interest via quantitative polymerase chain reaction (qPCR). Relative gene expression was calculated as 1.8^-ΔCt^ as described before [31], with ΔCt representing the difference between the Ct value of the gene of interest and the mean Ct values of the reference genes cleavage and polyadenylation specific factor 6 (*CPSF6*) and hypoxanthine phosphoribosyltransferase (*HPRT*). The quality of the qPCR products was evaluated by analyzing the melting curves and agarose gel electrophoresis.

### Immunohistochemistry

Day 28 samples were fixed in 4% formaldehyde (in PBS, pH 7.4)., dehydrated through a graded propane-2-ol series, and embedded in paraffin. From these, 5 μm thin microsections were generated. Following deparaffinization and rehydration, sections were treated with hyaluronidase (4 mg/mL in PBS, pH 5.5). Thereafter, sections were incubated with either pronase (1 mg/mL in PBS, pH 7.4, both from Roche Diagnostics, Mannheim, Germany) for type II collagen immunostaining, or with proteinase XXIV (0.02 mg/mL in PBS, pH 7.4, Sigma-Aldrich) for type X collagen or aggrecan immunostaining. For type X collagen immunostaining in hydrogels, pepsin (4 mg/mL in 0.01 M NaOH, Sigma-Aldrich) was used instead of proteinase XXIV.

Subsequently, sections were blocked with 5% bovine serum albumin. Human type II collagens, type X collagens, or aggrecans were visualized after incubation with a mouse anti-type II collagen antibody (1:1000, same as for Western blotting), a mouse anti-type X collagen antibody (1:1, as above), and a mouse anti-aggrecan antibody (1:25, clone HAG7D4 (7D4), #SM1353, Acris/OriGene, Rockville, MD, USA), respectively, and an ALP-coupled goat anti-mouse immunoglobulin G secondary antibody (ImmunoLogic, WellMed, MS Arnhem, The Netherlands) using the ImmPACT^®^ Vector^®^ Red Substrate kit (Vector Laboratories, Newark, CA, USA). Alkaline phosphatase activity in deparaffinized and rehydrated microsections was detected using NBT/BCIP substrate solution (in Tris-HCl (1M, pH 9.5); Merck).

### Statistical methods

The figure captions indicate the number of independent biological replicates for each experiment. Data was analyzed using SPSS Statistics (Version 29.0.0.0, IBM, Armonk, NY, USA). The Mann-Whitney U test was used to evaluate differences between groups. Box plots represent the interquartile range (IQR) extending between the 25^th^ and the 75^th^ percentiles (Q1, Q3), and lines inside the boxes represent the median. Whiskers extend to minimum and maximum values. Black circles in the graphs represent the single data points. Extreme outliers, indicated as empty circles, were determined using Tukey’s fences: Q3 + 3(Q3 – Q1). A paired Student’s *t* test was used for time-course studies. Results were considered statistically significant at p≤0.05.

## Results

### Endochondral commitment of MSCs in Heparin-PEG hydrogels in vitro

An initial pilot experiment indicated that MSC chondrogenesis in heparin-PEG hydrogels loaded with TGF-β was improved by additional TGF-β (10 ng/mL) supplementation of the chondrogenic medium (not shown). Next, we investigated whether these conditions induced a reproducibly strong deposition of cartilaginous matrix by MSCs. We prepared hydrogels with the same composition as in our previous study (60 µL PEG hydrogel formulation containing 22.4 mg/mL crosslinked heparin, 120 ng TGF-β1, 1.2×10^6^ MSCs) [18] and cultured them in standard TGF-β1-containing chondrogenic medium (10 ng/mL). Hydrogel-free standard pellets comprising 0.5×10^6^ MSCs without local TGF-β were used as controls. After four weeks, we detected substantial deposition of type II collagen and the main cartilage proteoglycan aggrecan in the hydrogels by histological analysis in all five independent experiments (Figure 1A). The cartilage matrix remained pericellular in only some areas, and not all cells had differentiated into chondrocytes, consistent with our previous in vivo results [18]. In control pellets and in the heparin group, strong and homogeneous deposition of type II collagen and aggrecan was detected and sufficient neocartilage was obtained to examine chondrocyte hypertrophy.

**Figure 1.**
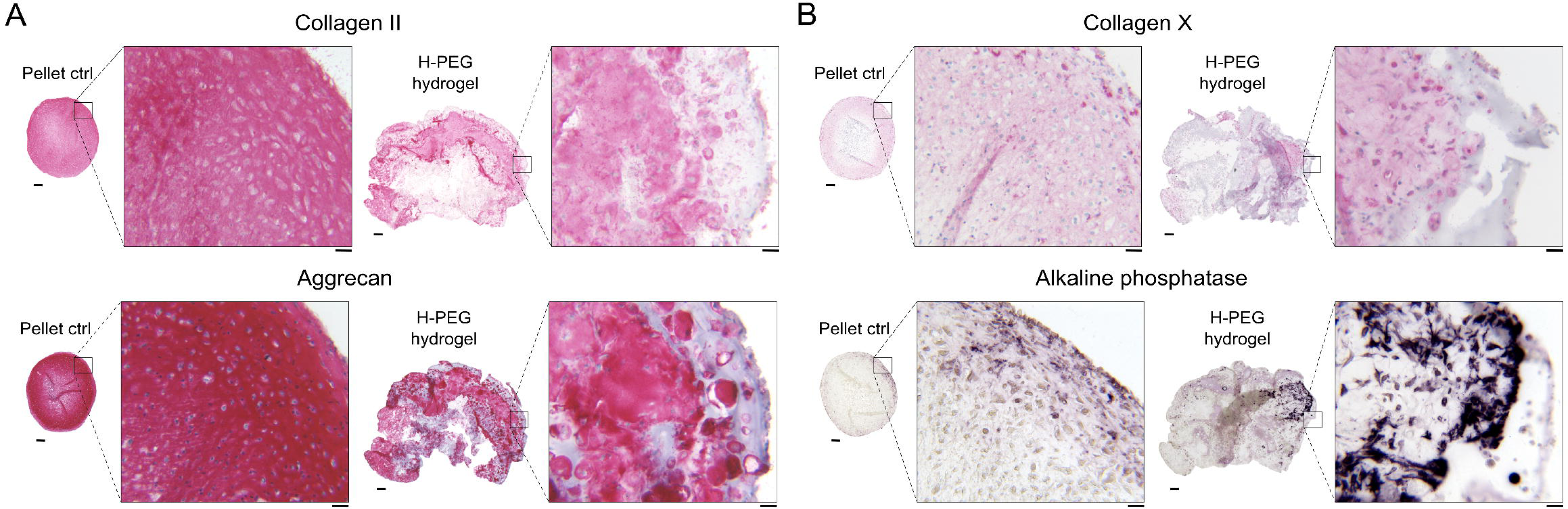
MSC in vitro chondrogenesis in heparin-PEG hydrogels versus standard carrier-free controls. MSCs were cultured as 3D pellets (pellet ctrl; 5×10^5^ cells) or in heparin-PEG hydrogels (H-PEG hydrogel; 1.2×10^6^ cells) containing 22.4 mg/mL crosslinked heparin and 120 ng TGF-β1 for 4 weeks in TGF-β1-containing (10 ng/mL) standard chondrogenic medium in vitro. Microsections of day 28 samples were assessed either via immunohistochemistry to detect **(A)** type II collagen or aggrecan, and **(B)** type X collagen, or by enzymatic activity staining to visualize alkaline phosphatase, as indicated (scale bar: overview = 200 µm, magnification = 50 µm). Results from one representative out of n = 5 experiments with independent MSC donor populations is shown.

We next addressed whether MSCs committed to chondral or endochondral differentiation in heparin-PEG hydrogels in vitro. Importantly, similar to the control pellets, the obtained neocartilage in the heparin-PEG group stained positive for type X collagen and the mineralizing enzyme ALP, indicating that chondrocytes became hypertrophic regardless of the presence of PEG-immobilized heparin when cultured in standard chondrogenic medium (Figure 1B). Of note, the MSC-derived neocartilage tissue which had formed in subcutaneous pouches in heparin-PEG hydrogels in our previous study was fully devoid of type X collagen and ALP [18], indicating that the lineage instructive ability of heparin-coupled TGF-β is context-dependent.

To further support our histological observations, we assessed gene expression levels of hypertrophy markers. Expression data was referred to collagen type II alpha 1 chain (*COL2A1*) gene expression to adjust for different chondrocyte quantities between pellet controls and hydrogels. In line with histological observations, qPCR analysis revealed strong expression of collagen type X alpha 1 chain (*COL10A1*) and *ALPL* mRNAs along with myocyte enhancer factor 2C (*MEF2C*) and *IBSP* in the heparin-PEG hydrogel group that was at least as high as in standard pellet controls (Figure 2, Supplementary Figure S1). Thus overall, MSCs underwent hypertrophic misdifferentiation in heparin-PEG hydrogels in standard chondrogenic medium, and heparin failed to redirect TGF-β-driven MSC chondrogenesis toward the desired chondral lineage. However, since modulating the heparin concentration in the hydrogel will inevitably change the mobility of TGF-β in the gel, its availability to the cells and thus the speed of chondrogenesis, the effect of heparin on cell signaling and chondrocyte hypertrophy was not addressed in this setup.

**Figure 2.**
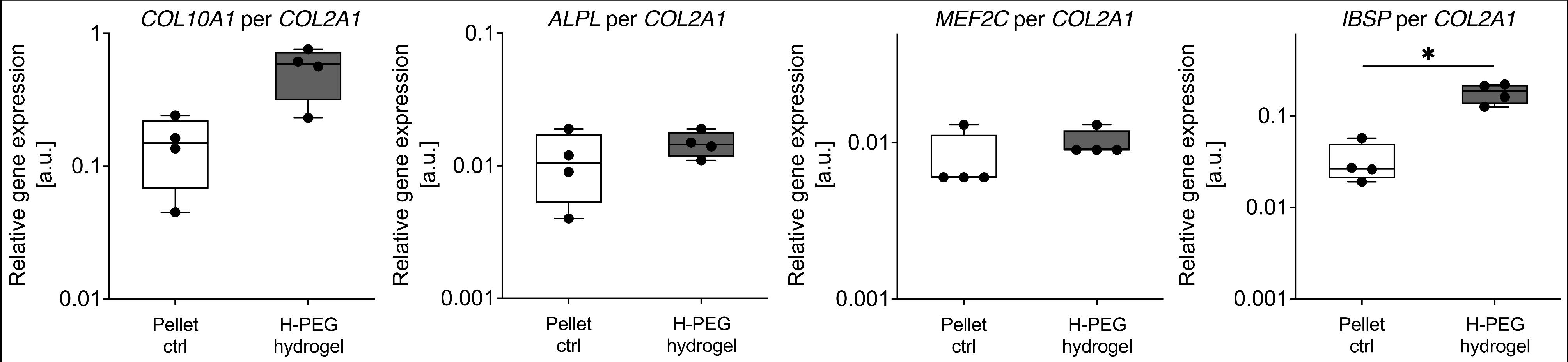
Gene expression analysis of MSCs cultured as 3D pellets or in heparin-PEG hydrogels. MSCs were cultured for 4 weeks in TGF-β1-containing (10 ng/mL) standard chondrogenic medium in vitro, either as 3D pellets (pellet ctrl; 5×10^5^ cells) or within heparin-PEG hydrogels (H-PEG hydrogel; 1.2×10^6^ cells; 22.4 mg/mL crosslinked heparin, 120 ng TGF-β1). The designated hypertrophy markers in day 28 samples are expressed as ratio to *COL2A1*, with *CPSF6* and *HPRT* used as reference genes. N = 4 experiments using independent MSC donor populations. Box plots were built as described in the statistics section. *p<0.05, Mann-Whitney U test.

### Effects of soluble heparin on lineage-directive cell signaling pathways

We next investigated the influence of soluble heparin on the activation of pro-chondrogenic and pro-hypertrophic signaling pathways at the initiation of chondrogenesis. Serum-free, defined standard chondrogenic medium with or without 10 ng/mL TGF-β1 and 6.25 μg/mL insulin was pre-incubated for 60 minutes with free soluble heparin at concentrations of 0, 10, 100, or 700 μg/mL and then used to stimulate pelleted passage 3 MSCs. As established [18], 700 μg/mL heparin was used as the maximum concentration. The 22.4 mg/mL heparin-containing PEG hydrogel formulation could not be tested in this experimental set-up due to its cytotoxic effects.

Western blotting showed that pro-chondrogenic SMAD2 phosphorylation was induced by TGF-β1 irrespectively of the presence of any tested heparin concentration (Figure 3A, Supplementary Figure S2A). By contrast, heparin dose-dependently decreased TGF-β1-induced pro-chondrogenic SMAD3 activation in all three examined donor populations (Figure 3B, Supplementary Figure S2B). A slight SMAD2/3 dichotomy was also observed in samples harvested after 28 days of chondrogenesis (Supplementary Figure S3A-B). Reduced SMAD3 activation under heparin treatment was in line with the previously observed reduction of TGF-β reporter activity in MSCs within heparin-PEG hydrogels compared to heparin-free controls [18]. Importantly, the SMAD1/9 phosphorylation which was induced by TGF-β1 at initiation of chondrogenesis was largely unaffected by 10 and 100 μg/mL heparin. It was, however, significantly decreased by 700 μg/mL heparin (Figure 3C, Supplementary Figure S2C). On day 28, no obvious reduction of TGF-β1-SMAD1/5/9 activation by any tested heparin concentration was observed (Supplementary Figure S3C). Remarkably, this is in contrast to suppressed pSMAD1/5/9 levels which were previously observed in vivo [18].

**Figure 3.**
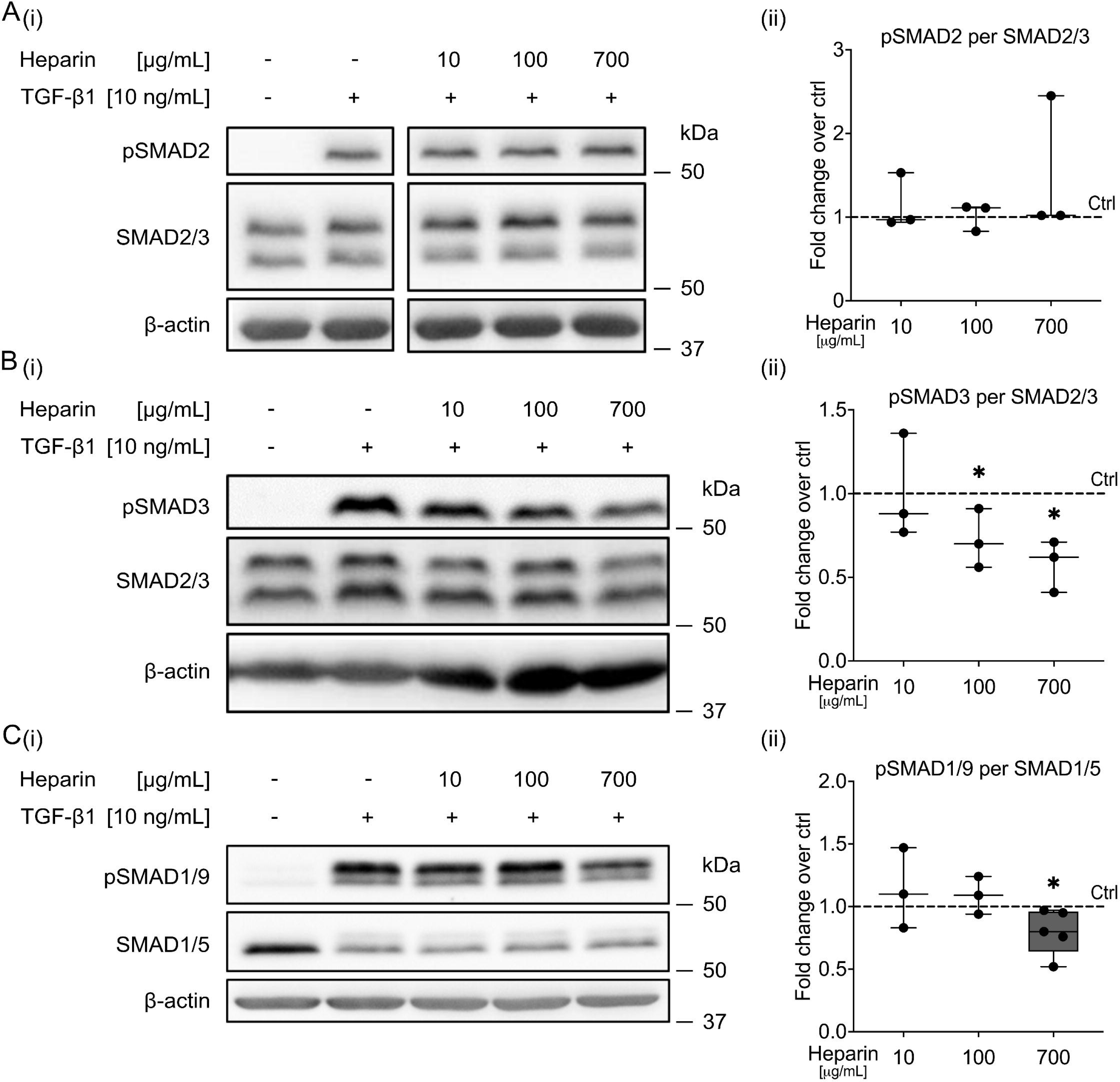
Effect of soluble heparin on TGF-β-induced SMAD activation in MSCs in vitro. Serum-free, defined chondrogenic medium, with or without TGF-β1 (10 ng/mL), was pre-incubated with soluble heparin (0, 10, 100, 700 µg/mL) for 60 minutes and then added to 5×10^5^ pelleted passage 3 MSCs. After 30 minutes, cells were harvested and whole protein lysates were prepared for Western blotting. **(A)** Phospho-SMAD2 and total SMAD2/3, **(B)** phospho-SMAD3 and total SMAD2/3, or **(C)** phospho-SMAD1/9 and total SMAD1/5 were detected. β-actin was used as an additional internal reference (all blots are shown in Supplementary Figure S2, uncropped blot pictures are provided as Supplementary Material S1). All samples in each line were run on the same gel and blotted onto one membrane. One out of n = 3-5 experiments using independent MSC donor populations is shown. Densitometric analysis of Western blots was normalized to total SMAD2/3 or total SMAD1/5, and control samples (+TGF-β1, no heparin) were each set to 1 (*p<0.05 vs. control, Mann-Whitney U test). Box plots were built as described in the statistics section.

Additionally, heparin supplementation mildly increased the insulin-induced phosphorylation of pro-chondrogenic AKT significantly (Figure 4A, Supplementary Figure S4A), which was previously shown to be important for TGF-β1-induced SMAD2 phosphorylation [4]. In turn, TGF-β1-stimulated pro-hypertrophic β-catenin accumulation was significantly reduced by all tested heparin concentrations (Figure 4B), which was in line with our previous in vivo observations.

**Figure 4.**
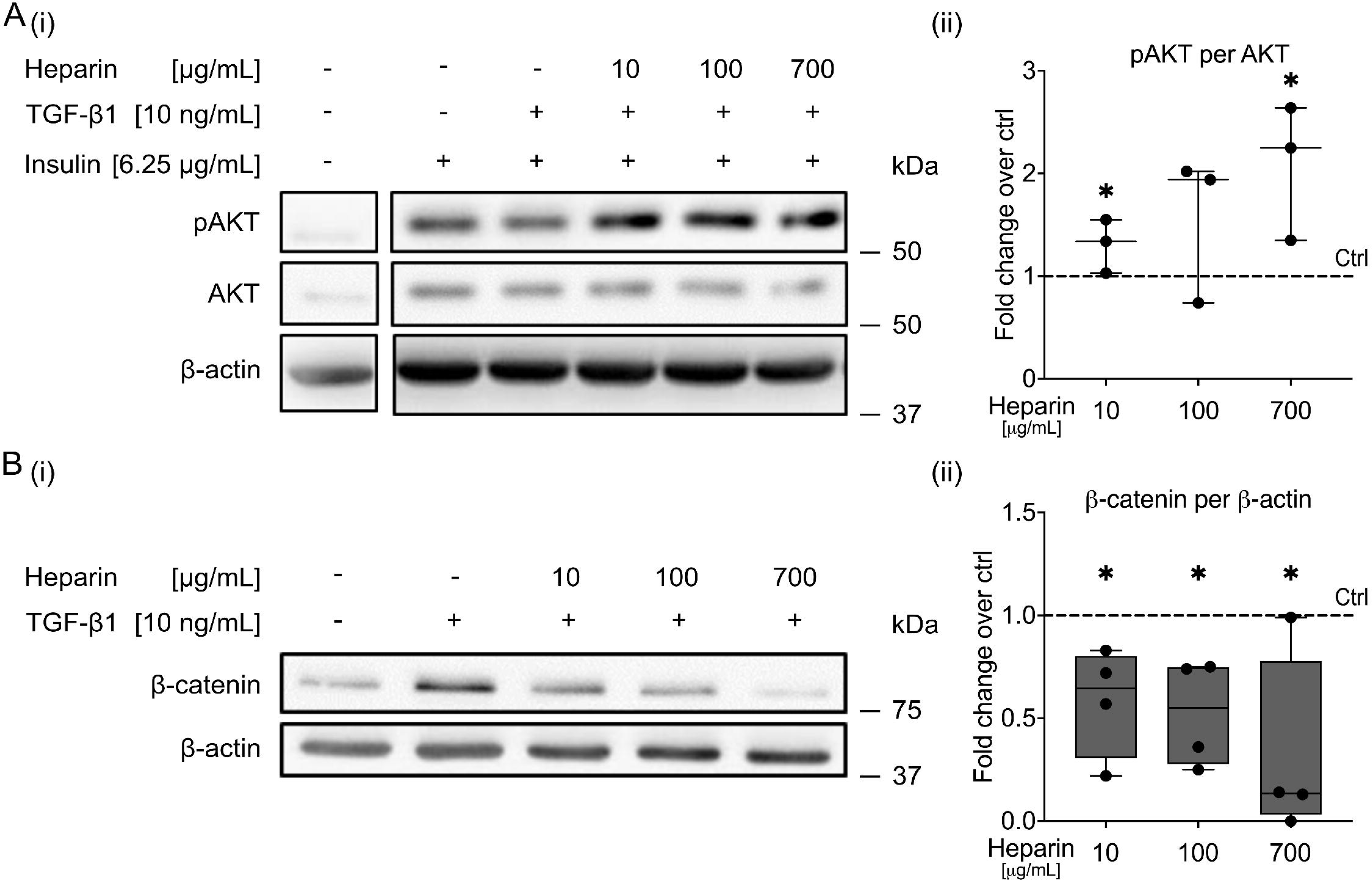
Effect of soluble heparin on AKT activation and β-catenin accumulation in MSCs in vitro. Serum-free, defined chondrogenic medium with or without TGF-β1 (10 ng/mL) and with or without insulin (6.25 µg/mL) was pre-incubated with soluble heparin (0, 10, 100, 700 µg/mL) for 60 minutes and then added to 5×10^5^ pelleted passage 3 MSCs. **(A)** After 30 minutes, samples were harvested and whole protein lysates were analyzed for phospho-AKT and total AKT using Western blotting. All samples in each line were run on the same gel and blotted onto one membrane. **(B)** Cytoplasmic-nuclear extracts of 3-hour samples were interrogated for β-catenin protein levels. β-actin was used as an internal reference. Representative blots of n = 3-4 independent experiments with independent MSCs are shown (all blots are shown in Supplementary Figure S4; uncropped blot pictures are provided as Supplementary Material S1). Box plots were built as described in the statistics section with a dashed line representing control samples set to 1 (+TGF-β1, +insulin, no heparin). *p<0.05 vs. control, Mann-Whitney U test.

Taken together, soluble heparin exhibited a surprising dichotomous effect on TGF-β/SMAD2/3 activation, inhibiting SMAD3 but not SMAD2 phosphorylation, which was possibly facilitated by enhanced insulin/AKT signaling. Concurrently, soluble heparin treatment reduced the activation of the endochondral WNT/β-catenin pathway but lowered SMAD1/5/9 activation only mildly and transiently. Together, these data suggested, that soluble heparin supplementation during MSC in vitro chondrogenesis may reduce the chondrogenic power and exhibit a mild anti-hypertrophic activity.

### Effect of soluble heparin on MSC in vitro chondrogenesis

We then investigated whether the observed maintained SMAD2 but reduced SMAD3 activation would reduce MSC in vitro differentiation into chondrocytes. Therefore, we determined gene expression levels of chondrocyte markers after 28 days of chondrogenic culture. While the expression of *COL2A1*, aggrecan (*ACAN*), and *SOX9* was strongly and significantly increased in the untreated control group compared to day 0 samples, as expected (Supplementary Figure S5A-C), this upregulation appeared reduced in samples cultured with 100 µg/mL or 700 µg/mL heparin. However, we observed a dose-dependent increase in median Ct values for all tested reference genes following treatment with heparin, and for the robust *CPSF6* and *HPRT* this became significant at 700 µg/mL (Supplementary Figure S5D). This observation indicated that soluble heparin treatment interfered with RNA isolation. Thus, the implications of gene expression analyses in the 100 and 700 µg/mL heparin groups remain unclear and should be interpreted cautiously.

We evaluated the neocartilage matrix accumulated during four weeks of chondrogenesis to determine whether the apparent reduction of *COL2A1*, *ACAN*, and *SOX9* gene expression on day 28 was evidence of reduced cartilage formation. Immunohistochemical and histological analyses revealed a strong and homogeneous deposition of type II collagen and aggrecan (Figure 5A). Cells showed the typical round chondrocyte morphology in all groups. However, the integrity of the extracellular matrix appeared compromised by 700 μg/mL heparin in all tested MSC donor samples and by 100 μg/mL heparin in 2 out of 5 MSC donors. ELISA, DMMB, and PicoGreen assays showed similar quantities of type II collagen, GAG/DNA, and DNA per pellet, respectively (Figure 5B-C, Supplementary Figure S5E), demonstrating similar levels of chondrogenic differentiation and matrix deposition among all groups.

**Figure 5.**
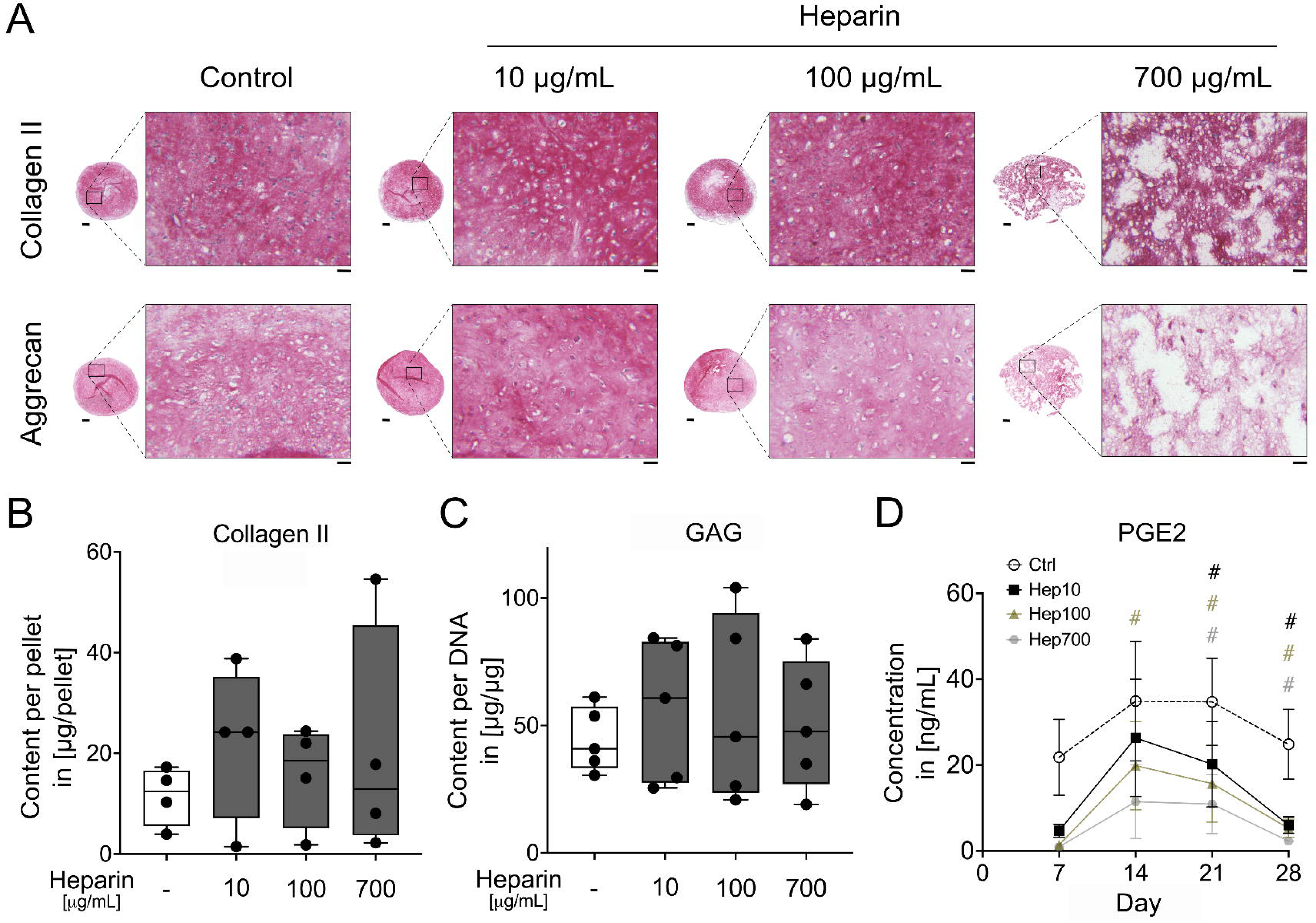
Effect of soluble heparin on MSC in vitro chondrogenesis. MSCs were cultured as 3D pellets for 28 days in standard chondrogenic medium (including 10 ng/mL TGF-β1 and 6.25 µg/mL insulin) supplemented with 0, 10, 100, or 700 µg/mL soluble heparin. **(A)** Paraffin microsections of day 28 samples were assessed for type II collagen and aggrecan via immunohistochemistry (scale bar: overview = 200 µm, magnification = 50 µm). Representative pictures of n = 5 experiments with independent MSCs are shown. **(B)** Type II collagen content per pellet was measured by ELISA (n = 4). **(C)** Proteoglycan content per pellet was assessed using DMMB assay and normalized to DNA amounts per pellet (n = 5). **(D)** Cell-conditioned medium was collected from 7-9 pellets at weekly intervals and analyzed for PGE2 levels using ELISA. Box plots were built as described in the statistics section and analyzed using Mann-Whitney U test. Line graphs show data as mean ± SEM. #p≤0.05 vs. control, paired Student’s *t* test.

Prostaglandin E2 (PGE2) was previously identified as an autocrine anti-hypertrophic mediator upregulated during MSC in vitro chondrogenesis [31]. ELISA revealed that PGE2 levels in cell culture supernatants were dose-dependently decreased by heparin throughout MSC chondrogenesis (Figure 5D), reaching –87±2% compared to the control group at culture termination (Supplementary Figure S5F). Adding heparin to PGE2 ELISA standards confirmed that heparin did not disturb PGE2 detection (Supplementary Figure S5G). Overall, while allowing strong chondrogenesis and cartilage formation at a low dose, increasing concentrations of soluble heparin compromised neocartilage integrity and strongly reduced levels of anti-hypertrophic PGE2. The high content of cartilaginous matrix in the pellets of all groups demonstrated maintained chondrogenic power despite reduced pSMAD3 levels, suggesting that the increased activation of AKT and sustained SMAD2 induction could fully compensate for the reduced SMAD3 induction.

### Soluble heparin is selectively anti-hypertrophic in vitro

Next, we assessed chondrocyte hypertrophy after 4 weeks of differentiation. As expected, the expression of the pro-hypertrophic transcription factor *MEF2C* along with *COL10A1*, *IHH*, the hedgehog target gene GLI family zinc finger 1 (*GLI1*), and *IBSP* mRNAs were significantly upregulated in control pellets (dashed line) compared to day 0 (Supplementary Figure S6A-F). Of note, treatment with 10 µg/mL heparin slightly but significantly reduced the day 28 levels of *MEF2C*, *IBSP,* and *ALPL* mRNAs, while the reduction of *COL10A1, IHH* and *GLI1* expression in all but one donors failed to reach significance (Figure 6A). The apparent dose-dependent reduction of these markers by higher heparin doses was inconclusive, as described above (Supplementary Figure S6A-G). Importantly, quantitative assessment of ALP enzyme activity in cell culture supernatants showed that all heparin concentrations strongly decreased secreted ALP levels during MSC chondrogenesis (Figure 6Bi). On day 28, the mean reduction reached dose-independently 85% in all groups compared to controls (Figure 6Bii). To rule out that the presence of heparin interfered with ALP detection, we added 10-700 µg/mL heparin to supernatants of heparin-free control pellet cultures. However, we found no effect on the detected enzyme activity (Supplemental Figure S6H). Importantly, Western blotting of pepsin-digested samples showed that type X collagen protein levels were not reproducibly reduced by heparin but rather strictly correlated with type II collagen protein levels (Figure 6C, Supplementary Figure S7). Of note, pepsin digestion is necessary to quantitatively release collagens from the strongly crosslinked cartilaginous matrix but also degrades all proteins that are commonly used as reference, including β-actin and glyceraldehyde-3-phosphate dehydrogenase (GAPDH).

**Figure 6.**
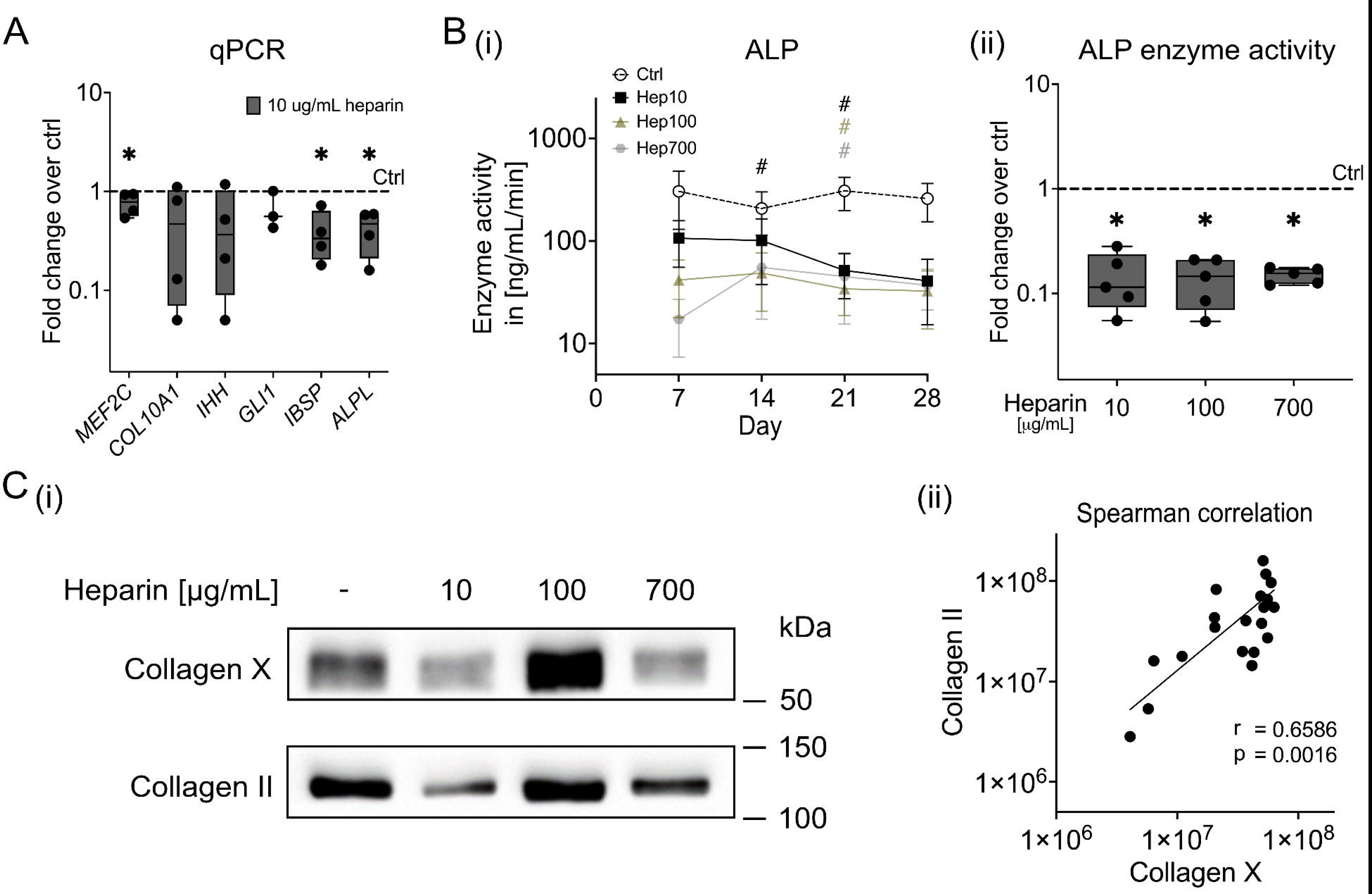
Effect of soluble heparin on hypertrophy during MSC chondrogenesis in vitro. MSC pellets were cultured for 4 weeks in standard chondrogenic medium (including 10 ng/mL TGF-β1 and 6.25 µg/mL insulin) containing soluble heparin (0, 10, 100, 700 µg/mL). **(A)** Gene expression levels of hypertrophy markers as designated, with *CPSF6* and *HPRT* used as reference genes (n = 3-4) and referred to untreated controls. **(B)** ALP enzyme activity was determined in the pooled supernatants of 7-9 MSC pellets at weekly intervals (n = 5). **(Ci)** Type X collagen content was determined via Western blotting in pepsin-digested day 28 MSC pellets, with type II collagen used as a reference. Representative blots of n = 5 from 4 independent donors are shown; all blots are depicted in Supplementary Figure S7. **(C_ii_)** Histogram of type X collagen versus type II collagen band intensities from Western blotting was analyzed by Spearman’s correlation, with the r-value and p-value indicated. Box plots were built as described in the statistics section. *p<0.05 vs. control, Mann-Whitney U test. Line graphs show data as mean + SEM. #p<0.05 vs. control, paired Student’s *t* test.

Collectively, our findings indicated that under standard in vitro chondrogenesis conditions neither PEG-immobilized heparin nor heparin in soluble form was capable of redirecting MSC fate into the chondral instead of the endochondral cell lineage. Yet, heparin was selectively anti-hypertrophic and strongly reduced the mineralizing enzyme ALP, albeit the important hypertrophy marker type X collagen remained unresponsive. Of note, this coincided with a WNT/β-catenin inhibition but ineffective silencing of TGF-β-mediated SMAD1/5/9 activation by heparin, as well as a reduction of anti-hypertrophic PGE2 levels throughout differentiation.

## Discussion

Preventing endochondral ossification and maintaining a stable chondral phenotype are key to achieving lasting, functional cartilage repair with MSC-based therapies. The currently most effective stabilizing strategies in vitro, namely PTHrP pulse treatment and WNT inhibition, are only selectively anti-hypertrophic and fail to inhibit type X collagen production [8, 17]. Conversely, in a subcutaneous mouse model, heparin-PEG hydrogels loaded with TGF-β silenced WNT– and SMAD1/5/9-pathways and permitted formation of permanent neocartilage from MSCs that was fully devoid of type X collagen and long-term resistant to calcification and bone formation [18]. The current in vitro study now demonstrates that critical, but so far unknown, contributions originating from the in vivo environment were mandatory to allow heparin-PEG to guide MSCs into the chondral instead of the endochondral lineage.

A thorough comprehension of the molecular mechanisms underlying the chondral versus endochondral lineage decision of MSCs is a prerequisite for controlling their directed differentiation both in fully defined conditions and after orthotopic implantation in the joint. For articular cartilage regeneration, in vitro chondrogenic pre-differentiation of MSCs is advantageous, as it allows for quality control and has shown significantly better histologic scores and morphologic characteristics of hyaline cartilage in large animal studies [1, 2]. Selecting sufficiently pre-differentiated grafts can mitigate the known variability of MSCs [32], thus ensuring reproducible therapy outcomes. Furthermore, the pre-formed cartilaginous extracellular matrix can increase the graft robustness and protect the cells from overload or toxic inflammation after implantation, thereby reducing failure rates in more challenging surgical settings. Directing MSCs towards a chondral instead of an endochondral cell fate during in vitro pre-differentiation is, therefore, a key requirement for the effective use of MSCs in future cartilage repair therapies.

For our initial investigations we chose the same heparin-PEG hydrogel formulation used in our prior in vivo experiments to address the ability of the biomaterial to instruct MSCs into the chondral lineage under defined standard TGF-β-driven in vitro chondrogenesis conditions. Since modulating heparin concentration in the hydrogel affects TGF-β mobility, availability and chondrogenesis speed, its impact on cell signaling and chondrocyte hypertrophy was not assessed in this setup. Switching to treatment with soluble heparin in vitro allowed us to evaluate the specific anti-hypertrophic effectivity of heparin, outside of in vivo microenvironmental influences.

We showed here for the first time that heparin reduced the activation of the pro-hypertrophic WNT/β-catenin signaling pathway and exhibited anti-hypertrophic activity during in vitro MSC chondrogenesis. Lowering the levels of the mineralizing enzyme ALP strongly (by 85%) already in its lowest applied concentration (10 µg/mL), heparin reached the effectiveness of PTHrP pulse treatment and WNT inhibition during chondrogenesis [8, 9, 17]. Also, its potency to reduce mean *IBSP* (−60%) and *IHH* expression (−51%) was within the range of previous results with PTHrP pulses and WNT inhibition [8]. Higher heparin concentrations of 100 µg/mL and beyond were, however, not advantageous because they compromised pellet integrity. Thus, we suggest using heparin in a 10 µg/mL dose as anti-hypertrophic intervention in future experiments. Notably, WNT inhibition and daily pulses with PTHrP require either the application of expensive proteins or may induce adverse effects due to solvents (e.g., DMSO); or they require complex and cumbersome application protocols. By contrast, heparin is cost-effective and can simply be applied with the routine medium (three times a week). Moreover, its extensive clinical application ensures access to high-quality grades and formulations; regulatory hurdles are well-mapped, and downstream translational applications are greatly facilitated.

One of the most important findings of this study was the observation of a dichotomous regulation of TGF-β-mediated SMAD2/3 activation in response to heparin treatment. While we observed SMAD3 regulation only at heparin concentrations higher than 10 µg/mL, it is likely that this phenomenon occurs over a broader concentration range, but was not detectable due to the limitations in sensitivity of Western blotting. The dichotomous response was surprising, since SMAD2 and SMAD3 are mostly reported to be coregulated. Consistent with this observation, our previous assays with the SB4 reporter showed a decreased SMAD3 reporter activity by TGF-β in MSCs cultured in the heparin-PEG hydrogels compared to heparin-free controls [18]. Obviously, heparin modulates the interaction of TGF-β with its receptors in a way that less SMAD3 is activated in vitro while SMAD1/5/9 activity is especially silenced in vivo. Since we previously demonstrated that insulin-AKT signaling enhances TGF-β-mediated SMAD2 phosphorylation during MSC chondrogenesis in vitro [4], we propose here that the stimulation of insulin-AKT activation by heparin supported SMAD2 activation by TGF-β. Importantly, reduced overall SMAD3 activation did not impair chondrogenic power according to unaffected amounts of extracellular matrix components in the neocartilage tissue. Thus, we propose that reduced SMAD3 activity can be tolerated when SMAD2 is sufficiently activated. This interpretation is consistent with the previous observation that genetic ablation of Smad2 affected mouse growth plate chondrocytes more severely than loss of Smad3, suggesting that Smad2 plays a more prominent role than Smad3 in chondrogenesis [33].

Yet like all other current anti-hypertrophic in vitro interventions, treatment with heparin remained only selectively effective and did not prevent MSCs from committing to endochondral development. Yet again, upregulation of the calcium-binding type X collagen that identifies the hypertrophic chondrocyte zone in the growth plate proved unresponsive to anti-hypertrophic treatment. By contrast, neocartilage that was fully devoid of type X collagen formed in heparin-PEG hydrogels loaded with TGF-β in vivo [18]. We previously proposed the balance between SMAD2/3 and SMAD1/5/9-mediated TGF-β signaling as one possible director of chondral versus endochondral lineage instruction of MSCs [9, 18], which then determines downstream regulation of endogenous pro-hypertrophic pathways (WNT, IHH). Consistent with this model, we found that SMAD1/5/9 activation was strongly suppressed in type X collagen-free chondral heparin-PEG explants [18] but remained largely intact in vitro, where MSCs committed to endochondral development despite the presence of heparin. Unfortunately, we currently lack effective tools to specifically inhibit SMAD1/5/9, as MSC chondrogenesis is incompatible with genetic engineering approaches. Nonetheless, our study provides additional important support that high SMAD1/5/9 activity plays a crucial role for instructing MSCs into the endochondral developmental lineage.

An important open question is why heparin used to retain TGF-β in a hydrogel can effectively instruct a chondral fate decision of MSCs in vivo but not in vitro. Given that high sulfation was required for chondral fate instruction and partial desulfation lead to endochondral differentiation, it appears likely that sulfation-dependent binding of bioactive factors to heparin in vivo plays a prominent role. Indeed, heparin is known to bind a multitude of cytokines including Gremlin [34], an enhancer of TGF-β-SMAD2/3 signaling and inhibitor of BMP-SMAD1/5/9 activation. Moreover, additional anti-hypertrophic signals in vivo are a likely possibility; PTHrP for example can be produced by normal subcutaneous adipose tissue [35]. Furthermore, the well-recognized anti-angiogenic activity of heparin [36] is also a likely contributor, as sufficient availability of calcium and blood supply are crucial for endochondral ossification and bone remodelling. Of note, we here showed that heparin reduced the endogenous production of PGE2 during MSC chondrogenesis, that we previously identified as an autocrine inhibitor of chondrocyte hypertrophy [31]. The standard chondrogenic medium contains high doses of the anti-inflammatory glucocorticoid dexamethasone that inhibits PGE2 synthesis [37]. Conversely, macrophages, endothelial cells and mast cells are ready sources of PGE2 in the subcutaneous niche and this may paracrinely contribute to suppress endochondral misdifferentiation of MSCs in heparin-PEG hydrogels in vivo. Altogether, while our data suggest in vivo available extracellular SMAD1/5/9 inhibitors and PGE2, further experiments are warranted to investigate the contribution of these and further potentially auxiliary bioactivities that aid TGF-β-loaded heparin-PEG hydrogels to instruct and stabilize the chondral lineage decision of MSCs in vivo.

One limitation of our study is, that we did not directly compare in vitro and in vivo chondrogenesis in this study but rather related to our previous results. Naturally, batch-related variations in the molecular weight distribution and maleimide modification of heparin may have occurred between studies, and their impact on the heparin potency remains unclear. Still, we observed stable chondral development of a subcutaneous implant with the heparin-PEG batch used in the current study (data not shown). A similar point can be made for comparing heparin treatment with other anti-hypertrophic interventions, specifically WNT inhibition and pulsed PTHrP treatment. While we focused on their shared inability to fully instruct MSCs into chondral development, a direct comparison would provide a more comprehensive understanding of their relative effectiveness.

## Conclusions

In essence, we showed here for the first time that heparin dichotomously inhibited SMAD3 but not SMAD2 activation by TGF-β without affecting the chondrogenic power of MSC cultures. Thus, like in the growth plate, SMAD2 appears to play a more prominent role than SMAD3 for in vitro chondrogenesis. We further propose that a stimulation of the insulin-induced AKT activation aids in maintaining phosphorylated SMAD2 levels. Moreover, we found that heparin exhibited an anti-hypertrophic activity that was comparable in selectivity and effectivity with that of WNT inhibition and pulsed PTHrP treatment, previously considered the most effective interventions in that regard, but offered advantages with respect to cost, technical simplicity and cell-compatible high-quality formulation. Most importantly, the in vitro effects of heparin fell short of its remarkable ability observed previously in vivo: retaining TGF-β and permitting formation of permanent neocartilage fully devoid of type X collagen and long-term resistant to endochondral ossification. While heparin appeared to sufficiently inhibit activation of the pro-hypertrophic WNT pathway, it had two notable limitations: incomplete silencing of SMAD1/5/9 activation and undesired inhibition of anti-hypertrophic PGE2. Thus, we here demonstrated that environmental contributions were essential to allow heparin-PEG to guide MSCs into the chondral instead of the endochondral lineage and their identification may be key to successful stabilization of MSC chondrogenesis also under fully defined in vitro conditions. Our ability to prevent endochondral ossification is crucial to pave the way for the safe and effective therapeutic use of MSCs for cartilage regeneration.

## Data availability

The authors confirm unrestricted access to all raw data, statistical analysis, and materials used in this study upon reasonable request.

## Abbreviations

ACAN: Aggrecan
ALP: Alkaline phosphatase
COL2A1: Collagen type II alpha 1 chain
COL10A1: Collagen type X alpha 1 chain
CPSF6: Cleavage and polyadenylation specific factor 6
GAPDH: Glyceraldehyde-3-phosphate dehydrogenase
GDF5: Growth and differentiation factor 5
ELISA: Enzyme-linked immunosorbent assay
GLI1: GLI family zinc finger 1
H-PEG: Heparin-PEG hydrogel
HPRT: Hypoxanthine phosphoribosyltransferase
IBSP: Integrin-binding sialoprotein
IHH: Indian Hedgehog
IQR: Interquartile range
MEF2C: Myocyte enhancer factor 2C
MMP: Matrix metallopeptidase
mRNA: Messenger ribonucleic acid
MSC: Mesenchymal stromal cell
PEG: Poly(ethylene glycol)
PGE2: Prostaglandin E2
PTHrP: Parathyroid-hormone related protein
qPCR: Quantitative polymerase chain reaction
sGAGs: Sulfated glycosaminoglycans
SMAD: Small mother against decapentaplegic
SDS: Sodium dodecyl sulfate
SOX9: SRY-box transcription factor 9
TGF-β: Transforming growth factor beta
WNT: Wingless-int

## Ethics declarations

### Ethics approval and consent to participate

All experiments of this study involving human material received approval from the Ethics Committee for Human Experimentation of the Medical Faculty of the Heidelberg University (S-452/2018, approval date: 06 August 2018; S-845/2019, approval date: 18 December 2019; S-554/2023, approval date: 11 October 2023), which acts in full compliance with the guidelines outlined in the latest version of the 1975 Helsinki Declaration. Human MSCs were isolated from bone marrow samples after obtaining written informed consent by the patients.

### Consent for publication

All authors approved the final manuscript to be published.

## CRediT authorship contribution statement

**Sven Schmidt:** Data curation, Formal analysis, Investigation, Methodology, Validation, Visualization, Writing – original draft, Writing – review & editing. **Safak Chasan:** Data curation, Formal analysis, Investigation, Methodology, Writing – review & editing. **Helen Dietmar:** Writing – original draft, Writing – review & editing. **Felicia Klampfleuthner:** Investigation, Methodology, Writing – review & editing. **Tilman Walker:** Resources, Writing – review & editing. **Uwe Freudenberg:** Resources, Writing – review & editing. **Wiltrud Richter:** Conceptualization, Funding acquisition, Project administration, Resources, Supervision, Writing – review & editing. **Solvig Diederichs:** Conceptualization, Formal analysis, Funding acquisition, Project administration, Resources, Supervision, Validation, Visualization, Writing – original draft, Writing – review & editing. All authors have read and agreed to the submitted version of the manuscript. **S. Schmidt** and **S. Diederichs** take full responsibility for the integrity of the work as a whole.

## Acknowledgments

The authors express their gratitude to Carina Binder for excellent technical assistance, the clinicians of the Orthopaedic University Hospital Heidelberg for supplying primary patient material, and the patients for the donation of bone marrow samples. The graphical abstract was created using Servier Medical Art which is licensed under CC BY 4.0 (https://smart.servier.com).

## Declaration of AI and AI-assisted technologies in the writing process

During the preparation of this work the authors used YoKI (meta-llama/Meta-Llama-3.1-70B) to assist with language editing and Perplexity (https://www.perplexity.ai) for literature research. After using these tools, the authors reviewed and edited the content as needed and take full responsibility for the content of the publication.

---

**Supplementary Figure S1.** QPCR analysis of MSCs cultured either as 3D pellets or in heparin-PEG hydrogels. In vitro culture of MSCs for 4 weeks in standard chondrogenic medium was performed either as 3D pellets (pellet ctrl; 5×10^5^ cells) or in heparin-PEG hydrogels (H-PEG hydrogel; 1.2×10^6^ cells; 22.4 mg/mL crosslinked heparin, 120 ng TGF-β1). **(A)** Day 28 Ct values for the indicated reference genes were determined by qPCR. **(B-C)** Baseline (day 0) and day 28 gene expression levels of the indicated chondrocyte and hypertrophy markers, with *CPSF6* and *HPRT* used as reference genes. N = 4 experiments using independent MSC donor populations. Box plots were built as described in the statistics section. *p<0.05 vs. day 0, Mann-Whitney U test.

**Supplementary Figure S2.** All Western blots included in Figure 3 quantifications. Detection of **(A)** phospho-SMAD2 and total SMAD2/3, **(B)** phospho-SMAD3 and total SMAD2/3, and **(C)** phospho-SMAD1/9 and total SMAD1/5 in whole protein lysates. β-actin served as an internal reference.

**Supplementary Figure S3.** Effect of soluble heparin on TGF-β-induced SMAD activation in MSCs in vitro. MSCs were cultured as 3D pellets for 4 weeks in standard chondrogenic medium supplemented with soluble heparin (0, 10, 100, 700 µg/mL). On day 28, serum-free, defined chondrogenic medium, with or without TGF-β1 (10 ng/mL), was pre-incubated with soluble heparin (0, 10, 100, 700 µg/mL) for 60 minutes and then added to the MSC pellets. After 30 minutes, cells were harvested and whole protein lysates were prepared for Western blotting. **(A)** Phospho-SMAD2 and total SMAD2/3, **(B)** phospho-SMAD3 and total SMAD2/3, or **(C)** phospho-SMAD1/9 and total SMAD1/5 were detected (n = 1). β-actin was used as an additional internal reference. All samples in each line were run on the same gel and blotted onto one membrane.

**Supplementary Figure S4.** All Western blots included in Figure 4 quantifications**. (A)** Whole protein lysates were used to detect phospho-AKT and total AKT. **(B)** β-catenin was assessed in the cytoplasmic-nuclear fraction of whole protein lysates. β-actin was used as an internal reference. The transmembrane protein CD81 was used to confirm the successful separation of the cytoplasmic-nuclear fraction from the membrane fraction.

**Supplementary Figure S5.** MSC in vitro chondrogenesis in the presence of soluble heparin. MSCs were cultured as 3D pellets for 4 weeks in standard chondrogenic medium (including 10 ng/mL TGF-β1 and 6.25 µg/mL insulin) that was supplemented with soluble heparin (0, 10, 100, 700 µg/mL). **(A-C)** Gene expression levels of chondrocyte markers as designated, using *CPSF6* and *HPRT* as reference genes (n = 5). **(D)** Day 28 cycle threshold (Ct) values for the indicated reference genes were assessed by qPCR. **(E)** DNA content per pellet was assessed using PicoGreen fluorescent probe. **(F)** Differentiating MSCs on day 28 were analyzed for PGE2 secretion levels using ELISA. **(G)** The PGE2 standard solution from the PGE2 immunoassay kit was supplemented with 700 µg/mL soluble heparin. PGE2 levels were assessed via spectrophotometry (n = 1). Box plots were built as described in the statistics section with a dashed line representing control samples set to 1. *p<0.05 vs. day 28 control samples, Mann-Whitney U test.

**Supplementary Figure S6.** Hypertrophic development of MSC-derived chondrocytes in presence of soluble heparin. MSCs were cultured as 3D pellets for 4 weeks in standard chondrogenic medium supplemented with soluble heparin (0, 10, 100, 700 µg/mL). **(A-G)** Gene expression levels of hypertrophy markers as designated, using *CPSF6* and *HPRT* as reference genes (n = 5). **(H)** Heparin was added to MSC-conditioned positive control supernatants as specified. ALP enzyme activity in the culture supernatants was assessed spectrophotometrically via substrate conversion (n = 2). Box plots were built as described in the statistics section. Extreme outliers according to Tukey’s Fences test are indicated with empty circles. *p<0.05 vs. day 28 control samples, Mann-Whitney U test.

**Supplementary Figure S7.** Western blot analysis of type X collagen in all MSC donor populations included in Figure 6. Type II collagen was used as an internal reference (uncropped blot pictures are provided as Supplementary Material S1).

**Supplementary Table S1.** Alphabetical list of primer sequences utilized for qPCR analysis.

