## Supplementary figures and images for "Dichotomous SMAD2/3 regulation and selective anti-hypertrophic activity of heparin during in vitro chondrogenesis of mesenchymal stromal cells"

### Supplementary Figure S1

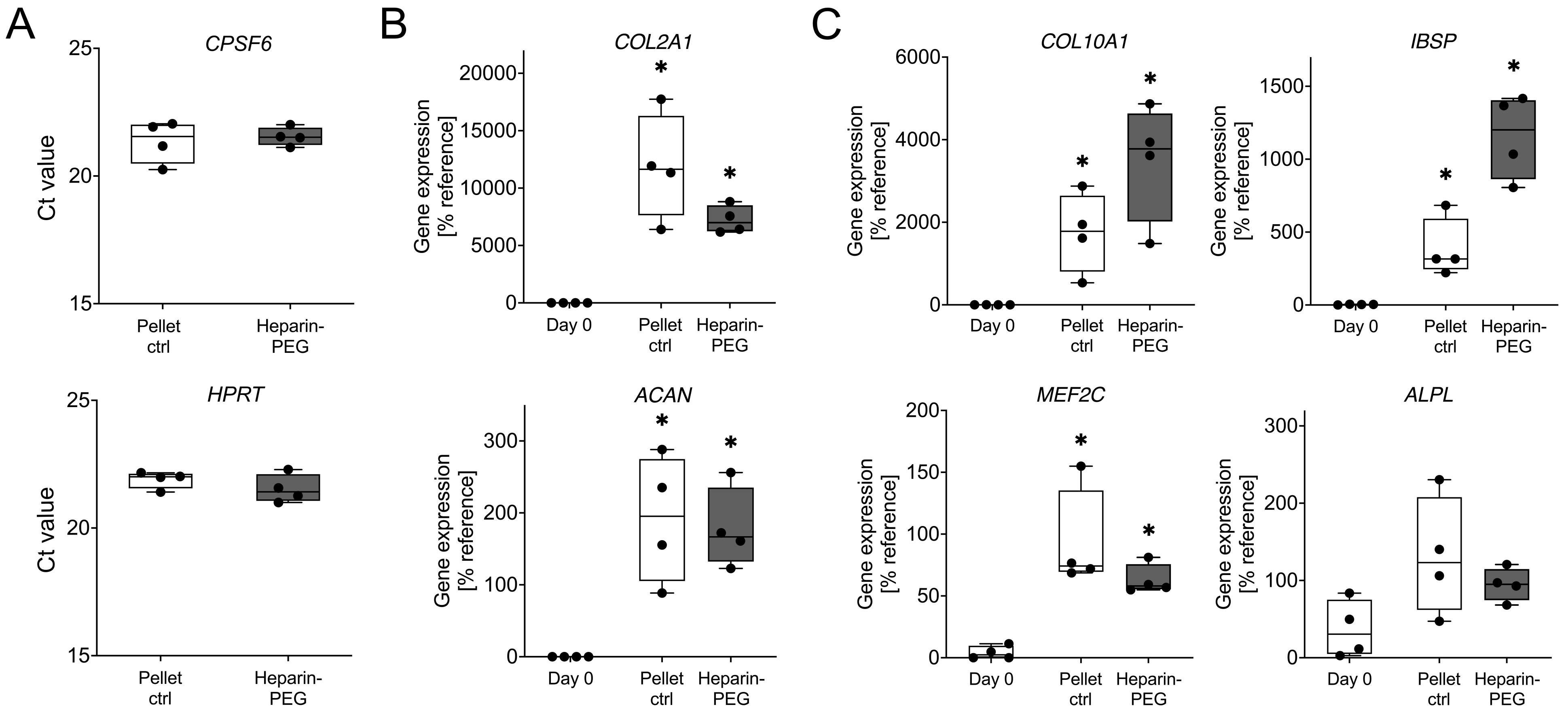

### Supplementary Figure S2

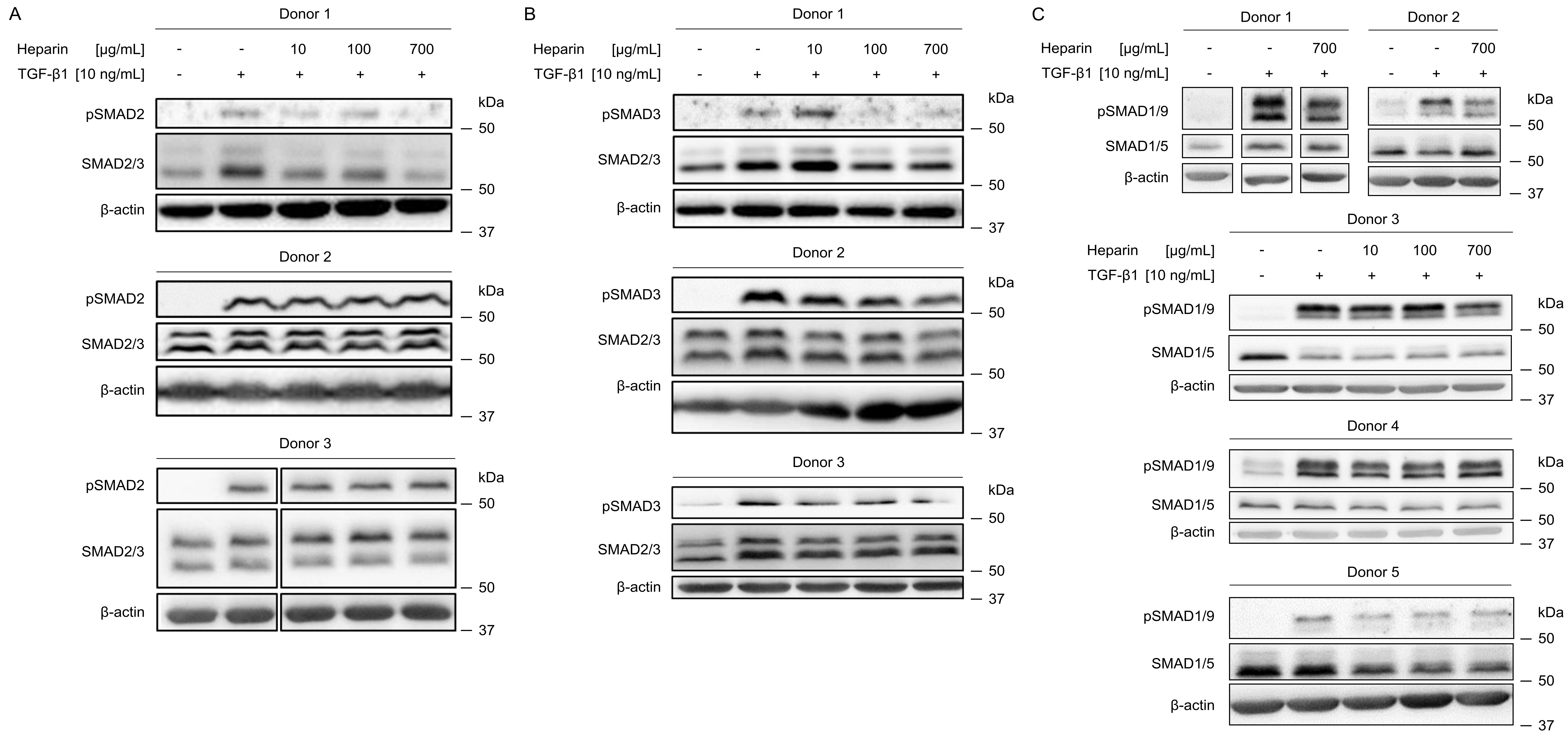

### Supplementary Figure S3

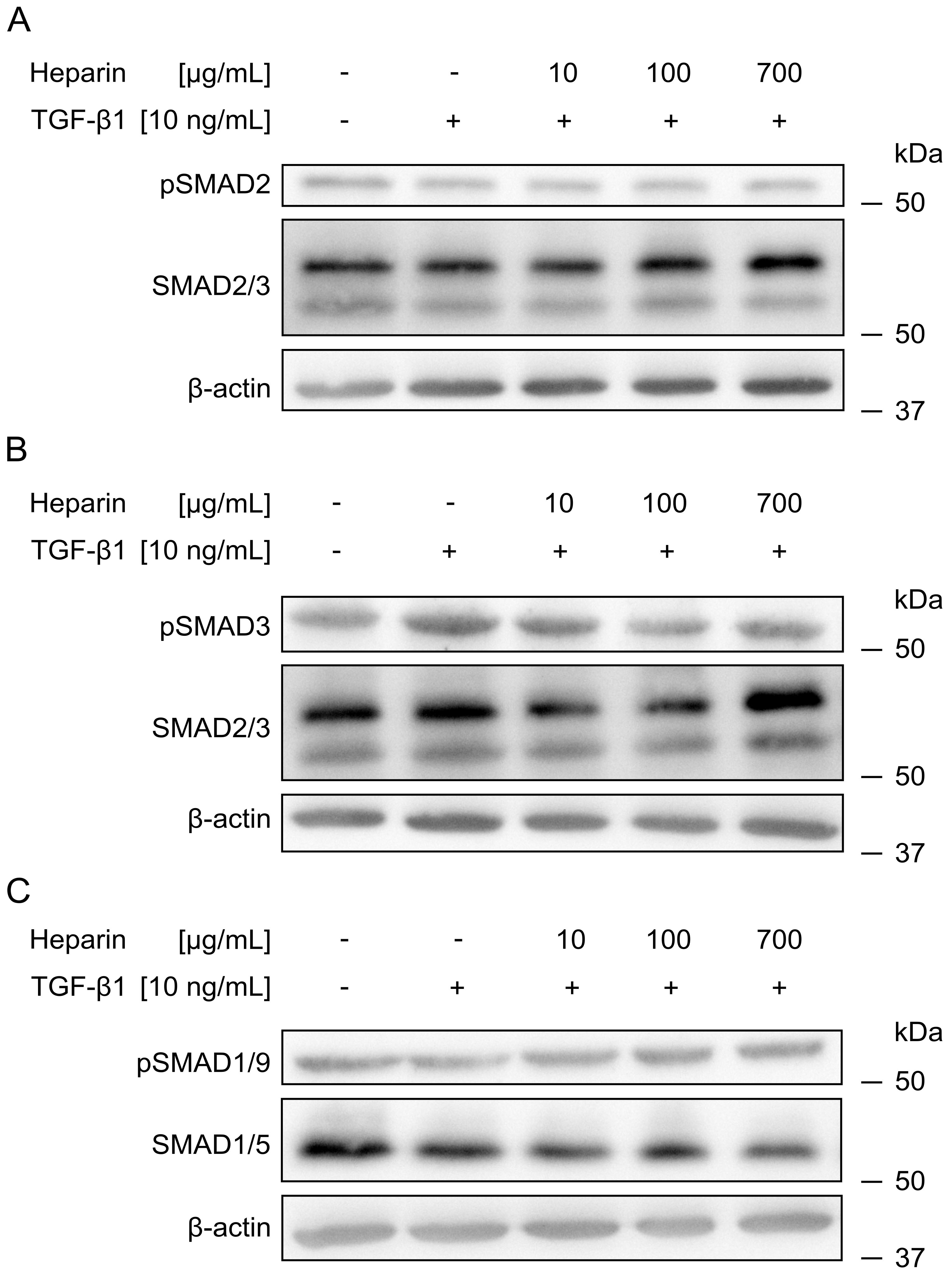

### Supplementary Figure S4

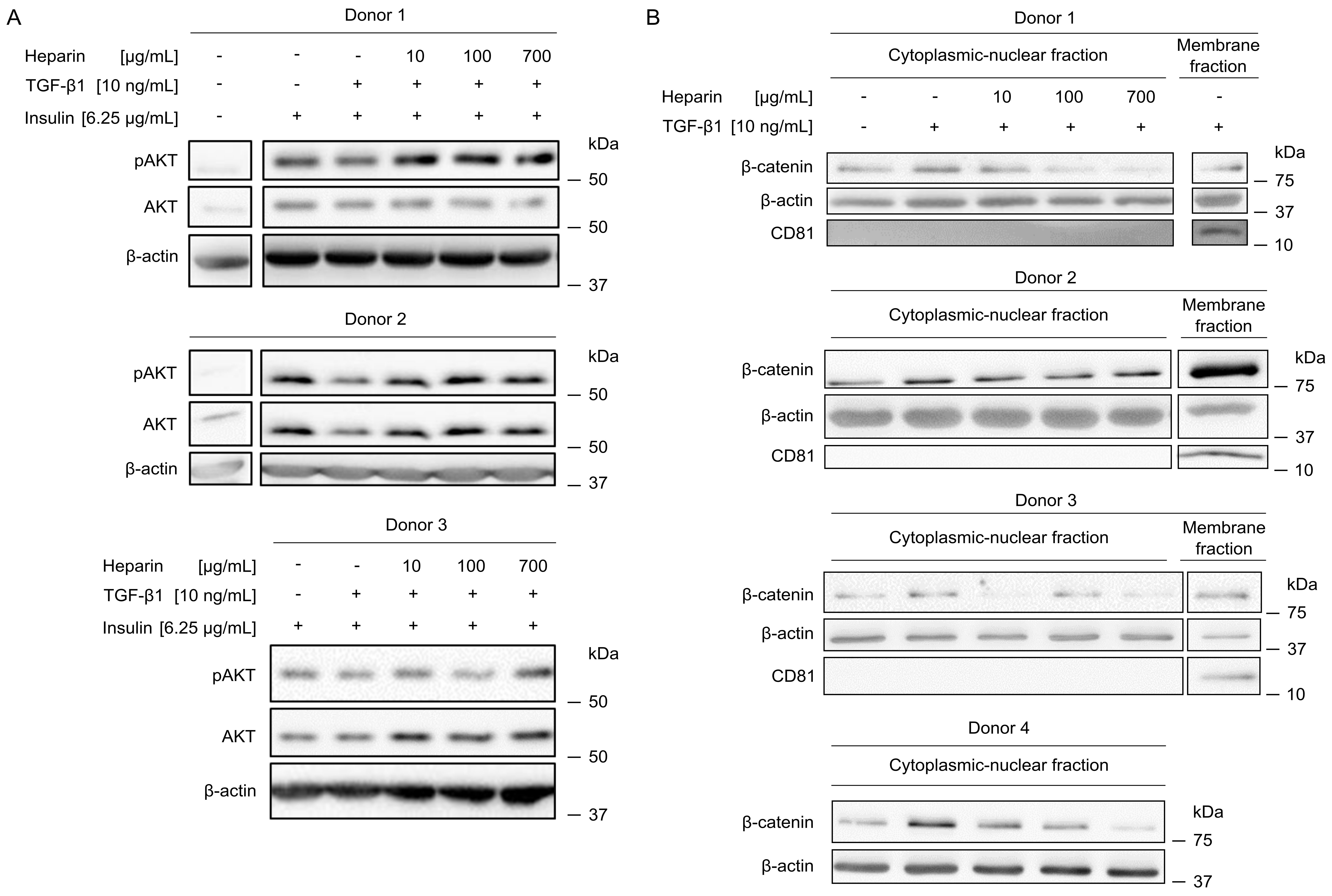

### Supplementary Figure S5

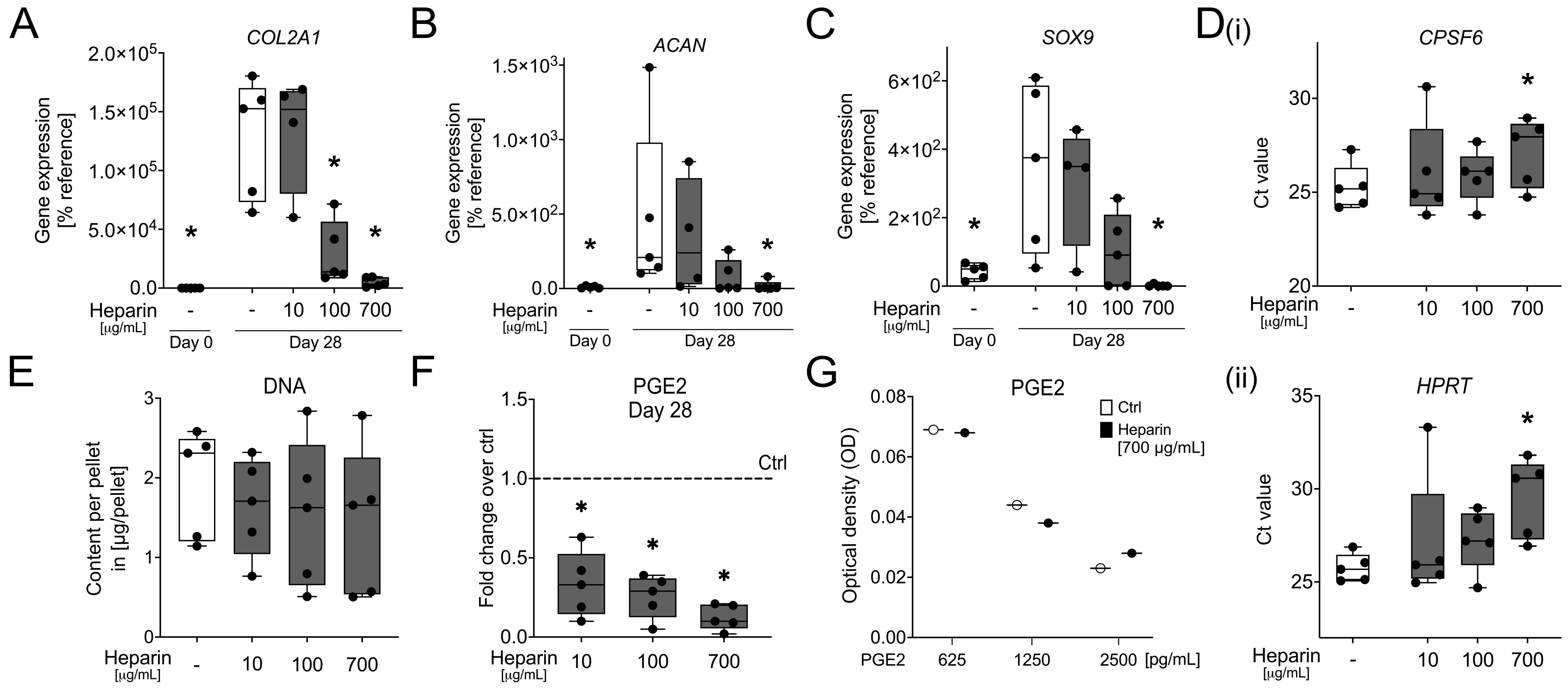

### Supplementary Figure S6

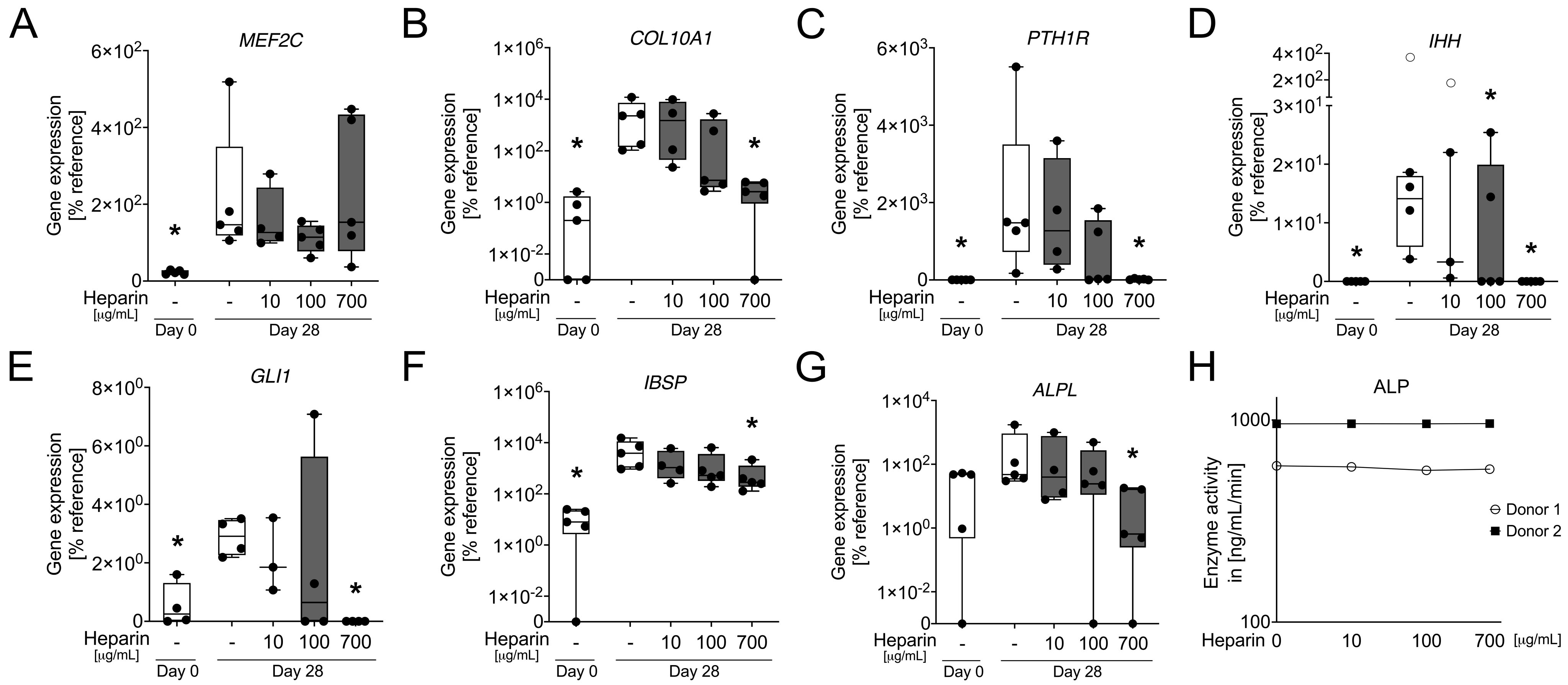

### Supplementary Figure S7

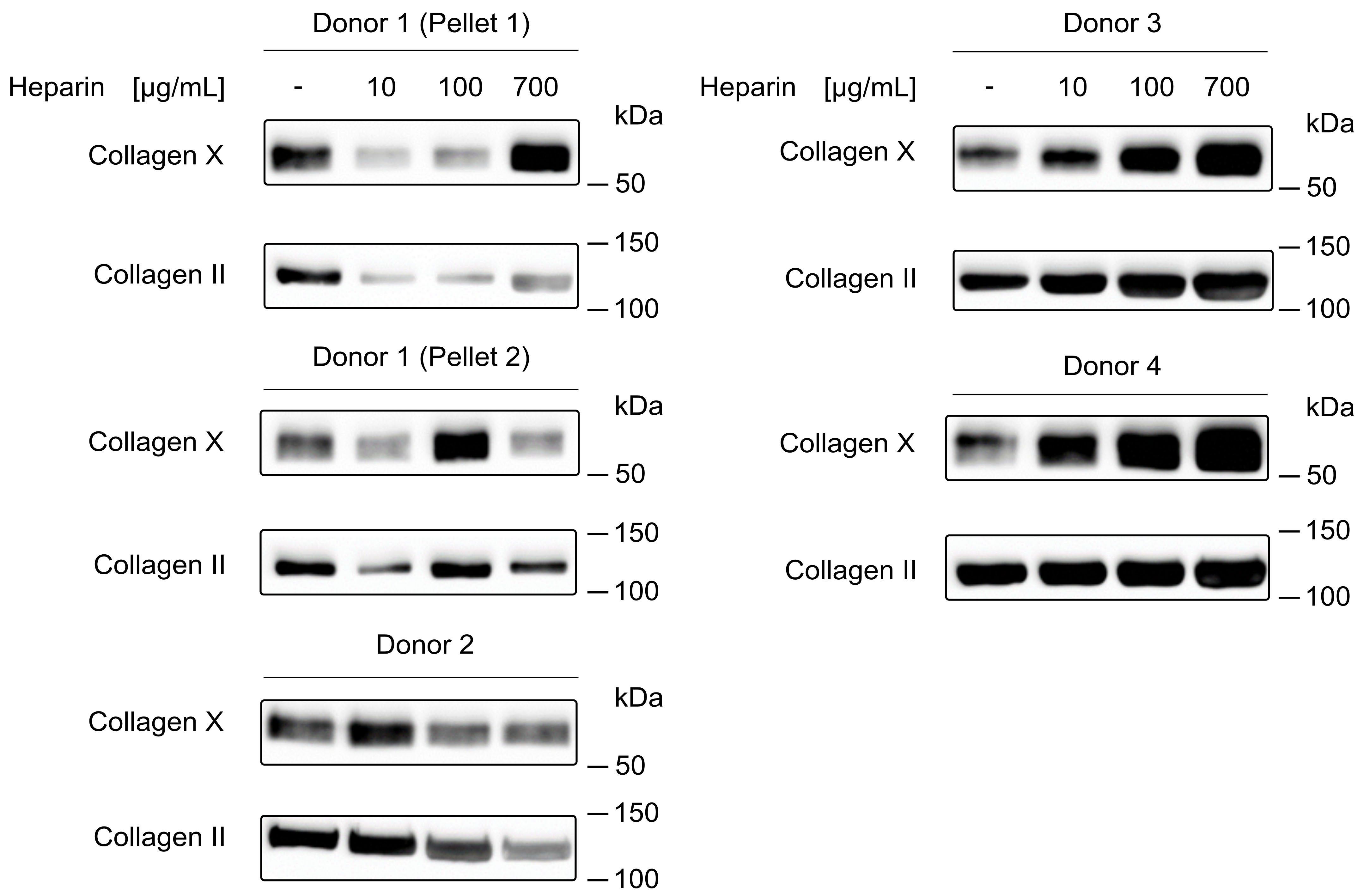
