## Supplementary Material S1 for "Dichotomous SMAD2/3 regulation and selective anti-hypertrophic activity of heparin during in vitro chondrogenesis of mesenchymal stromal cells"

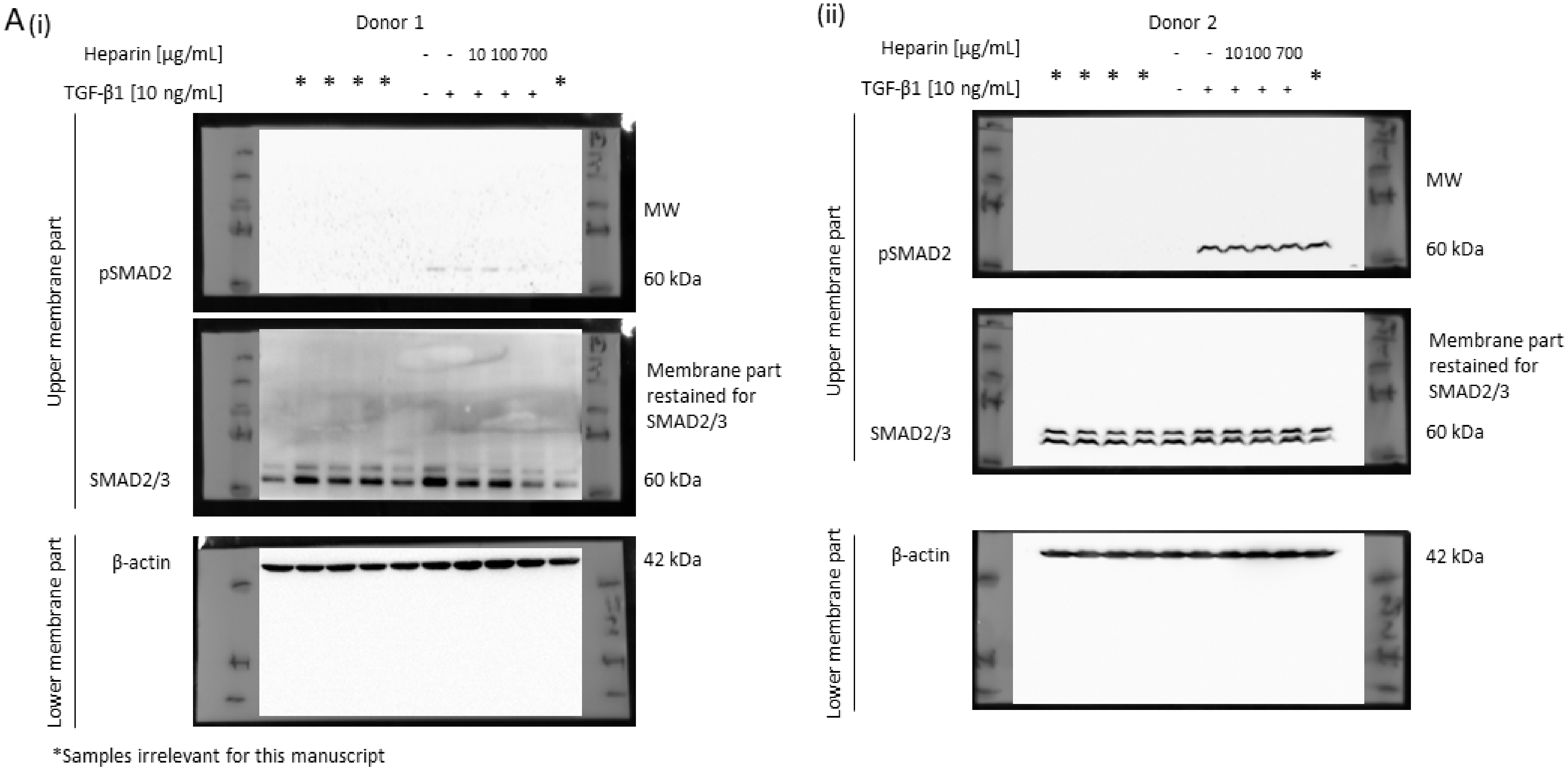


**Supplementary Material S1.** Western blots included in this study shown as full uncropped images.

(iii)

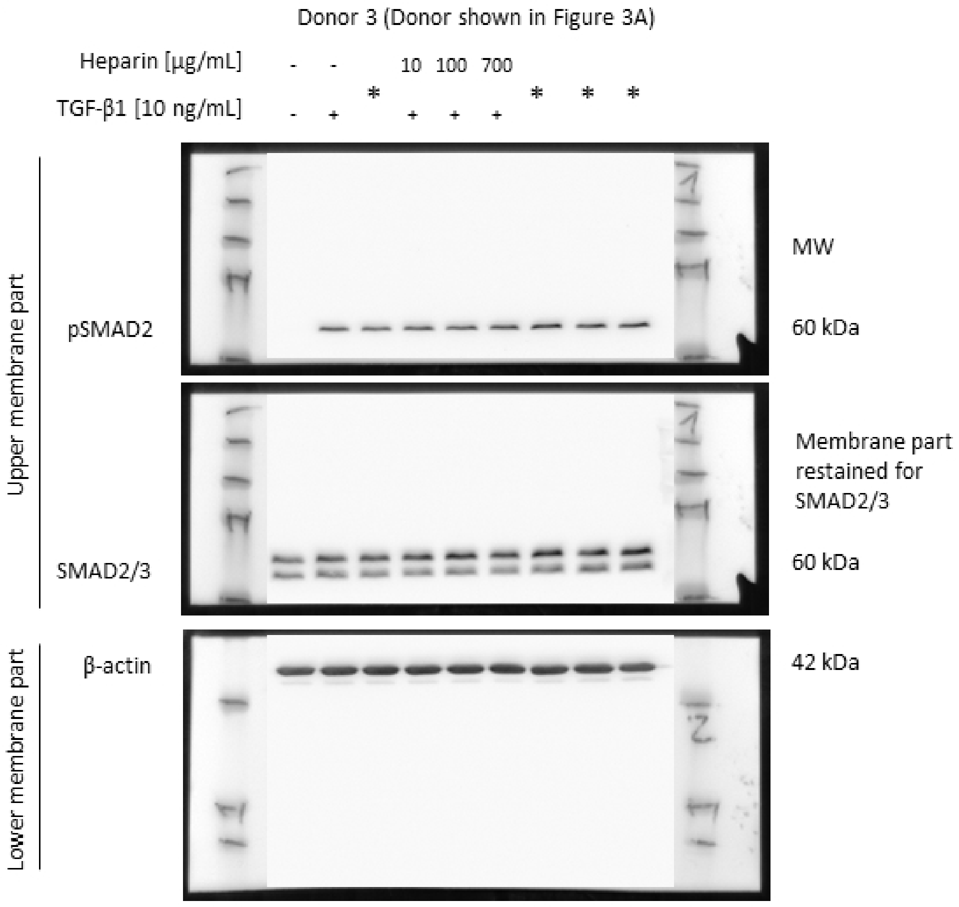


*Samples irrelevant for this manuscript

**Supplementary Material S1.** Western blots included in this study shown as full uncropped images.

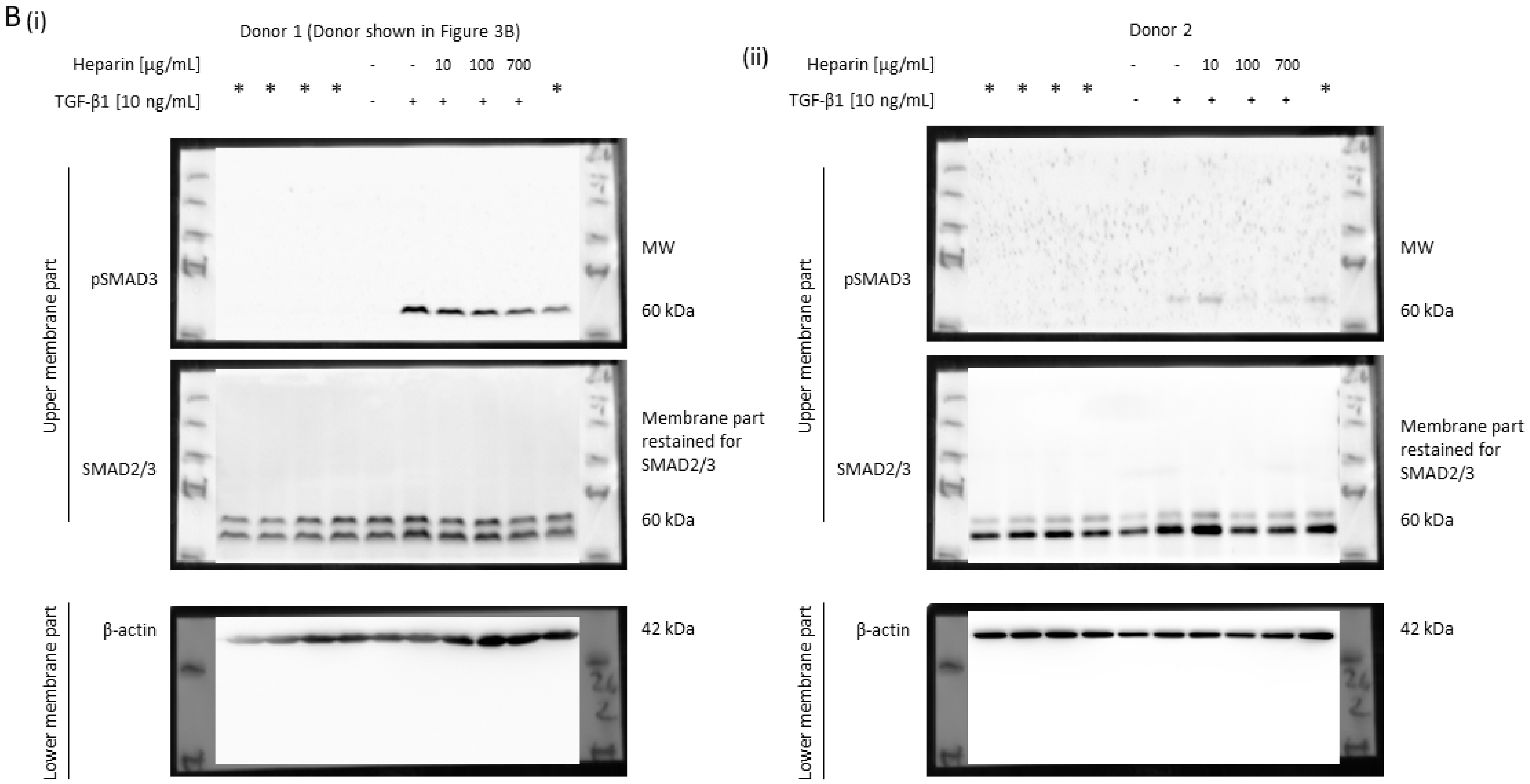


*Samples irrelevant for this manuscript

**Supplementary Material S1.** Western blots included in this study shown as full uncropped images.

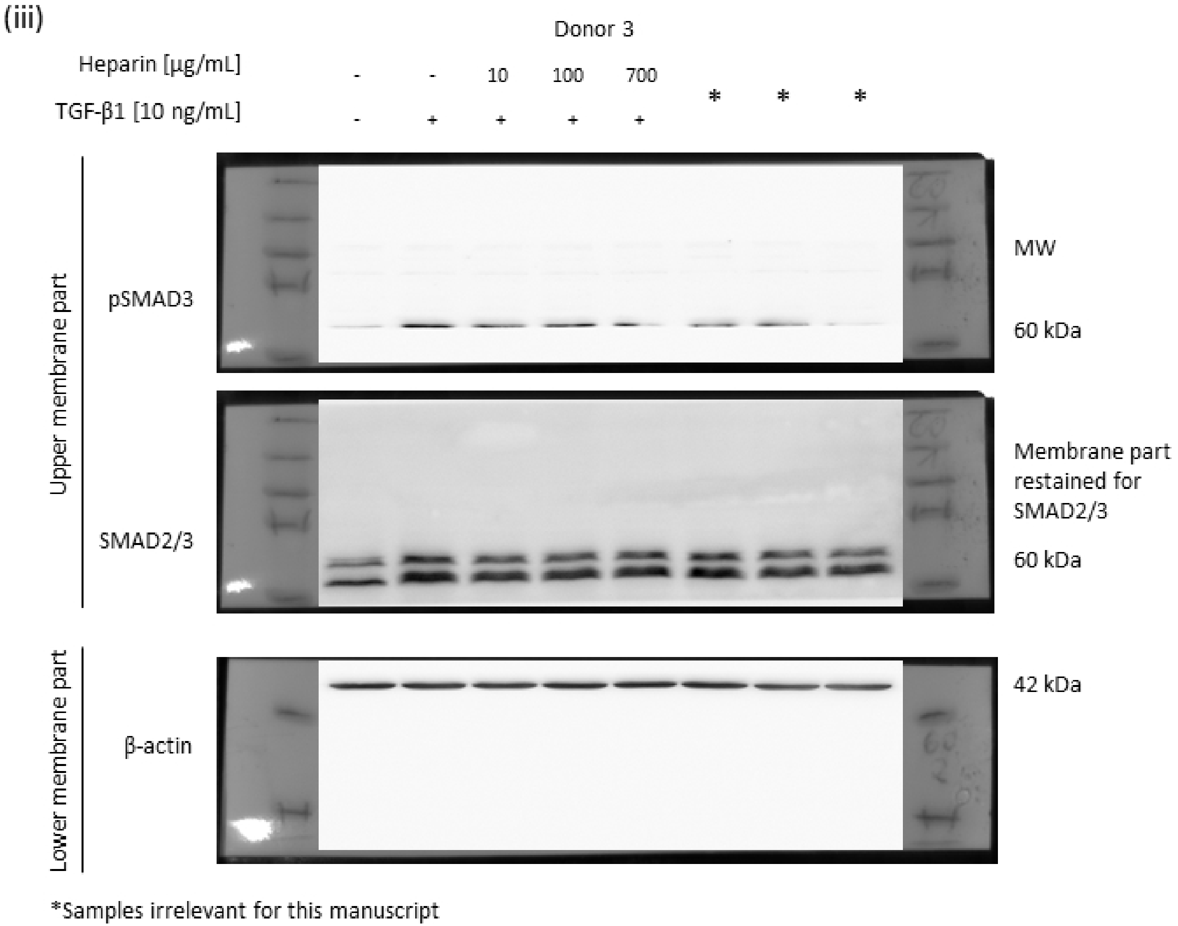


**Supplementary Material S1.** Western blots included in this study shown as full uncropped images.

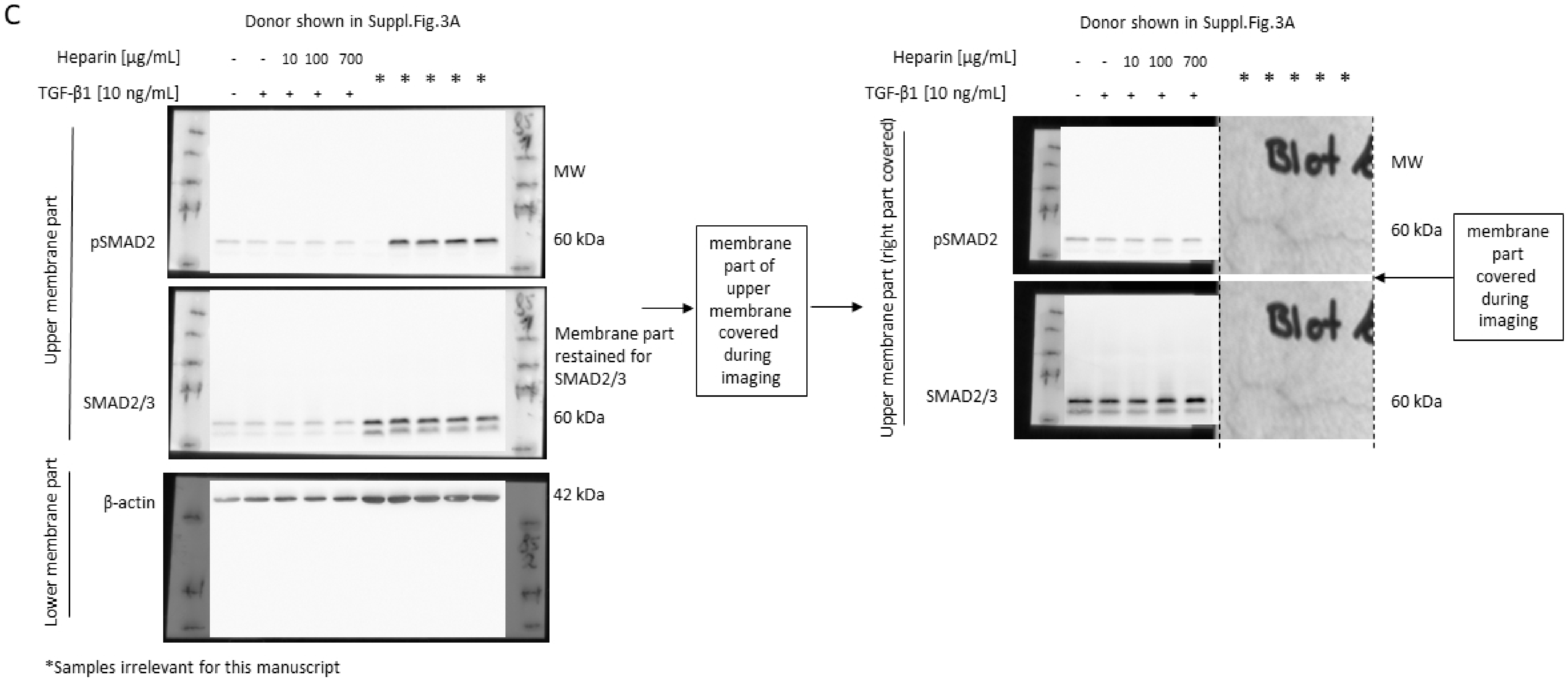


**Supplementary Material S1.** Western blots included in this study shown as full uncropped images.

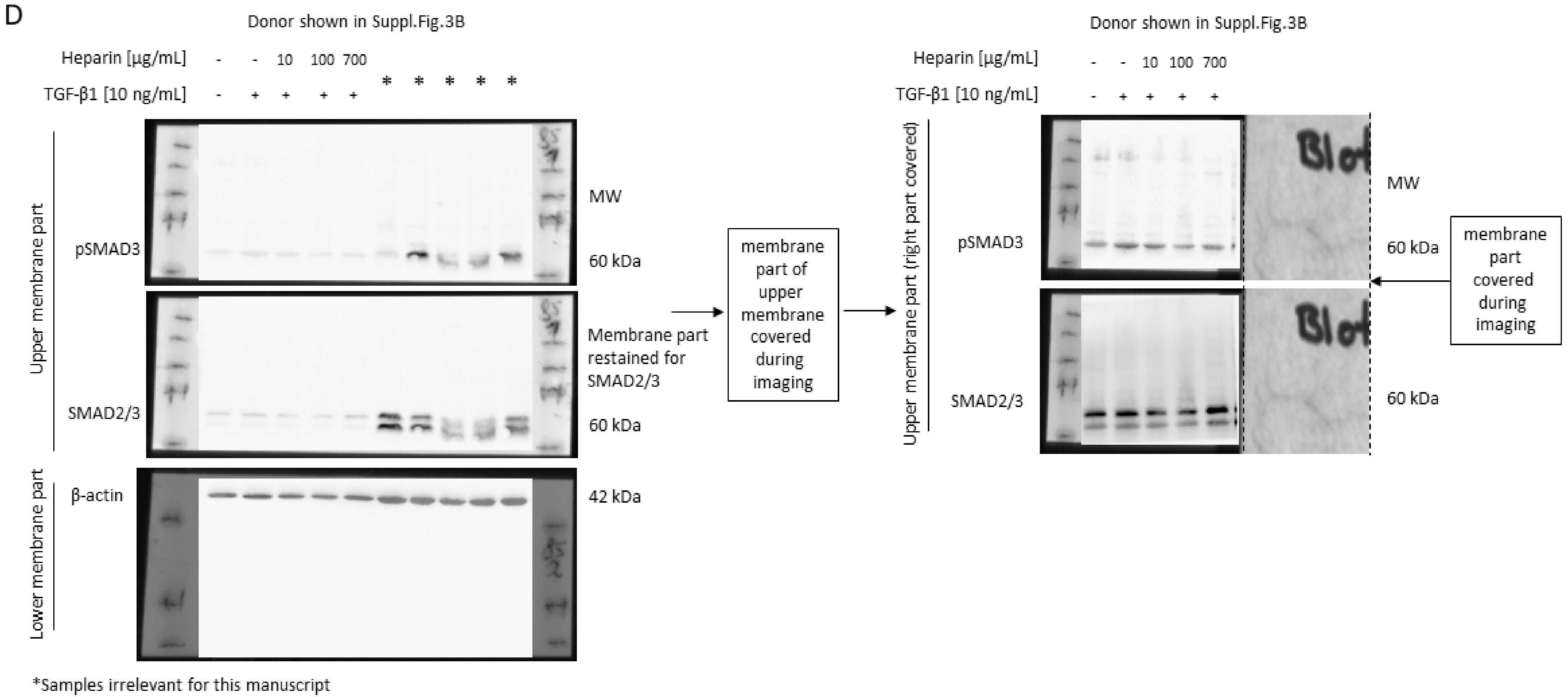


**Supplementary Material S1.** Western blots included in this study shown as full uncropped images.

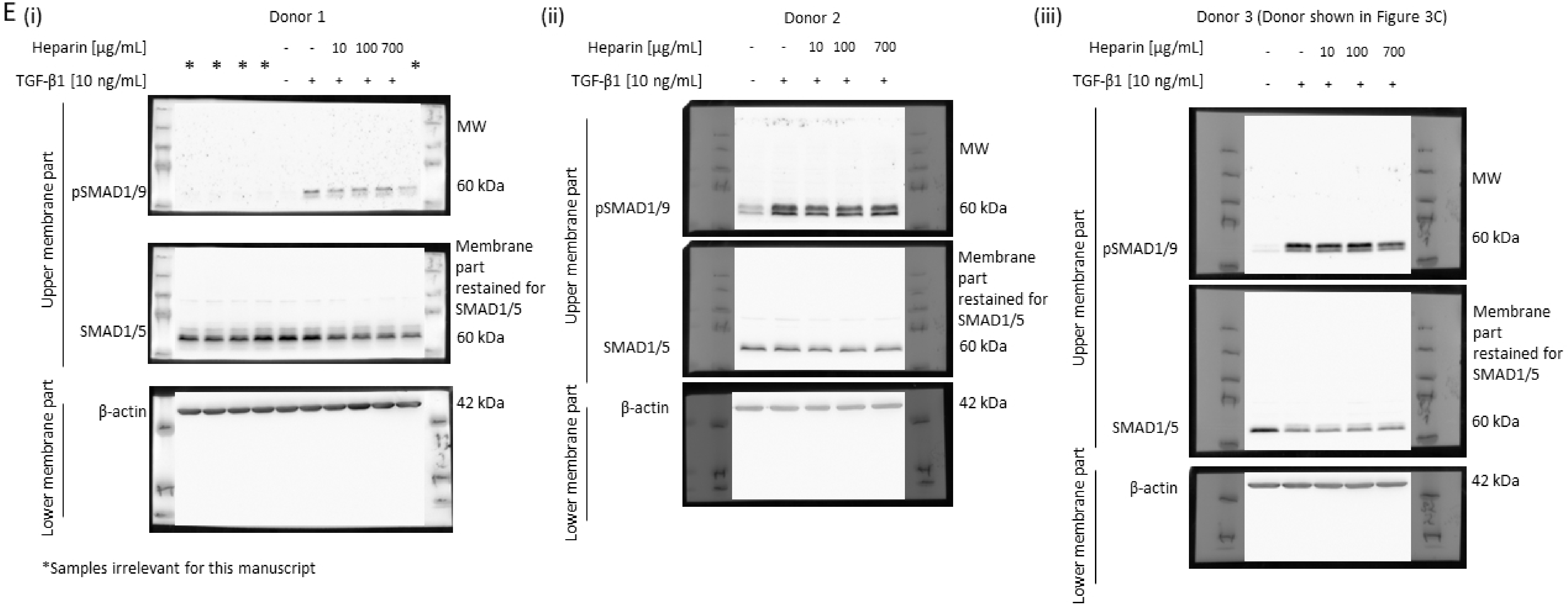


**Supplementary Material S1.** Western blots included in this study shown as full uncropped images.

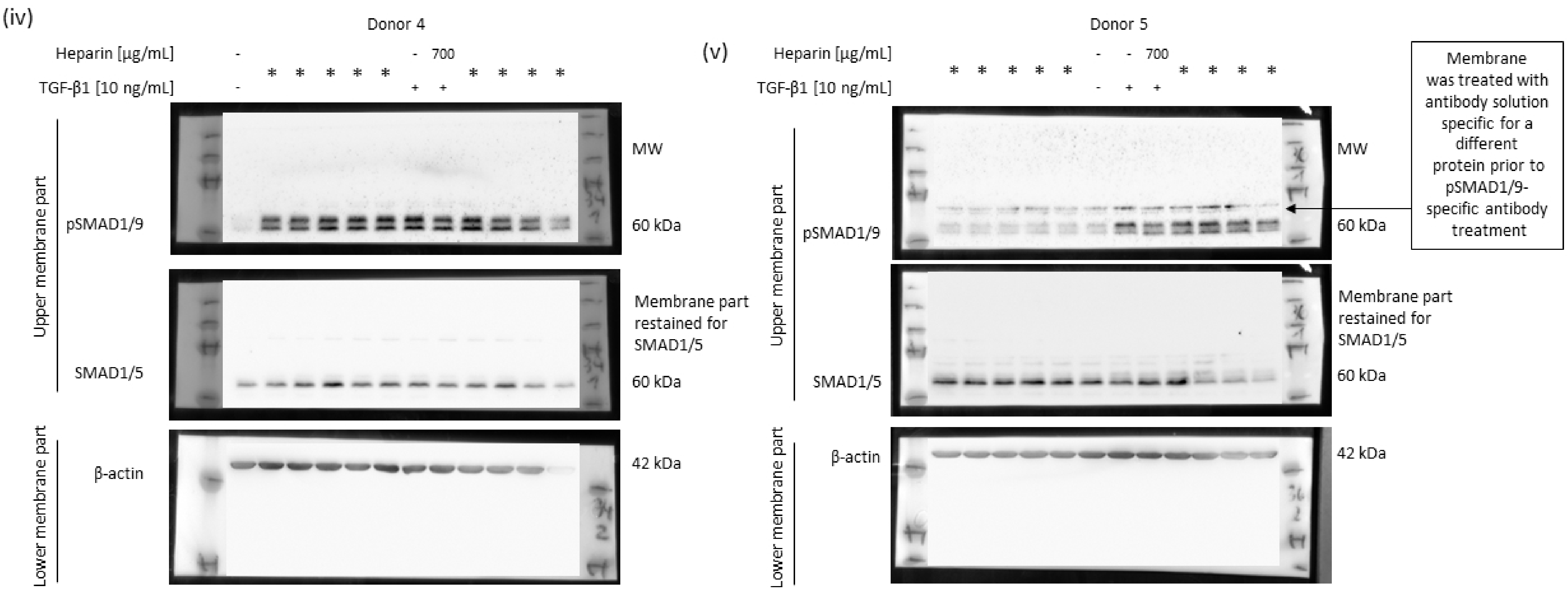


*Samples irrelevant for this manuscript

**Supplementary Material S1.** Western blots included in this study shown as full uncropped images.

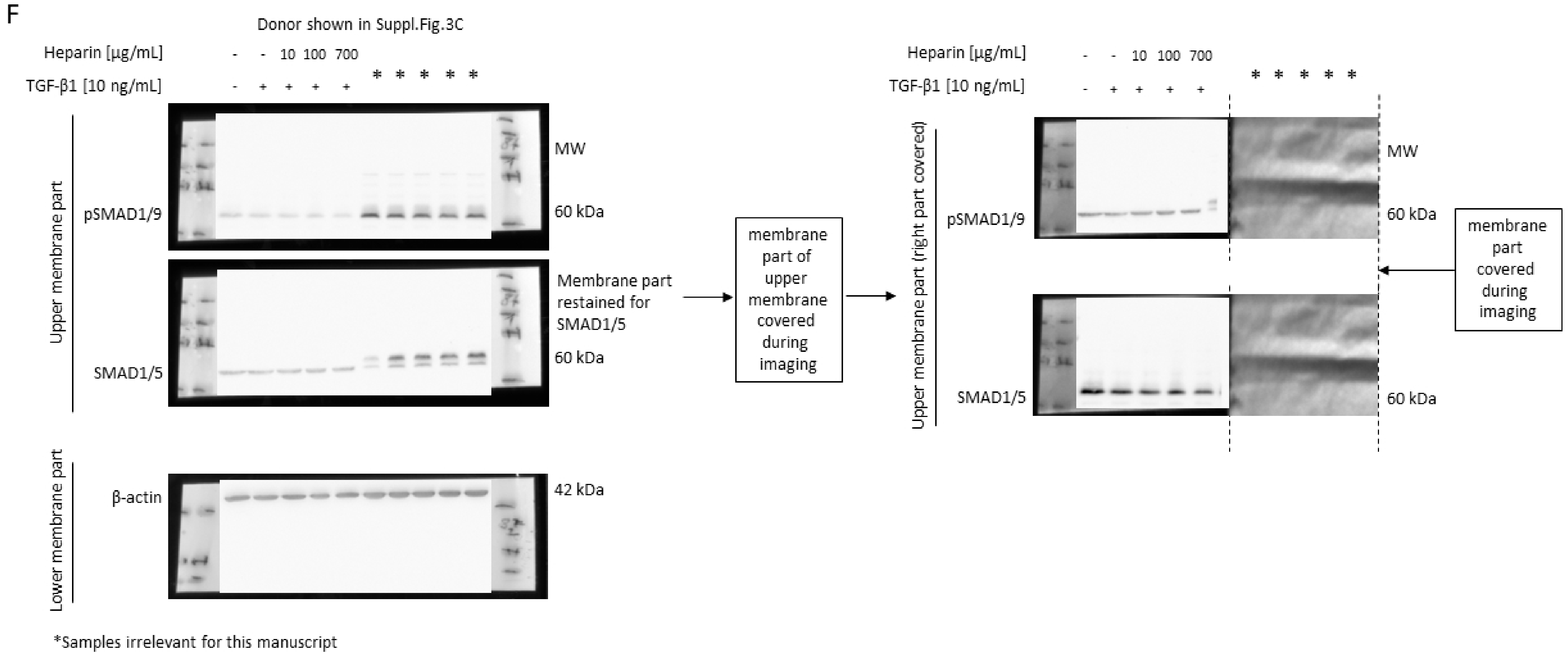


**Supplementary Material S1.** Western blots included in this study shown as full uncropped images.

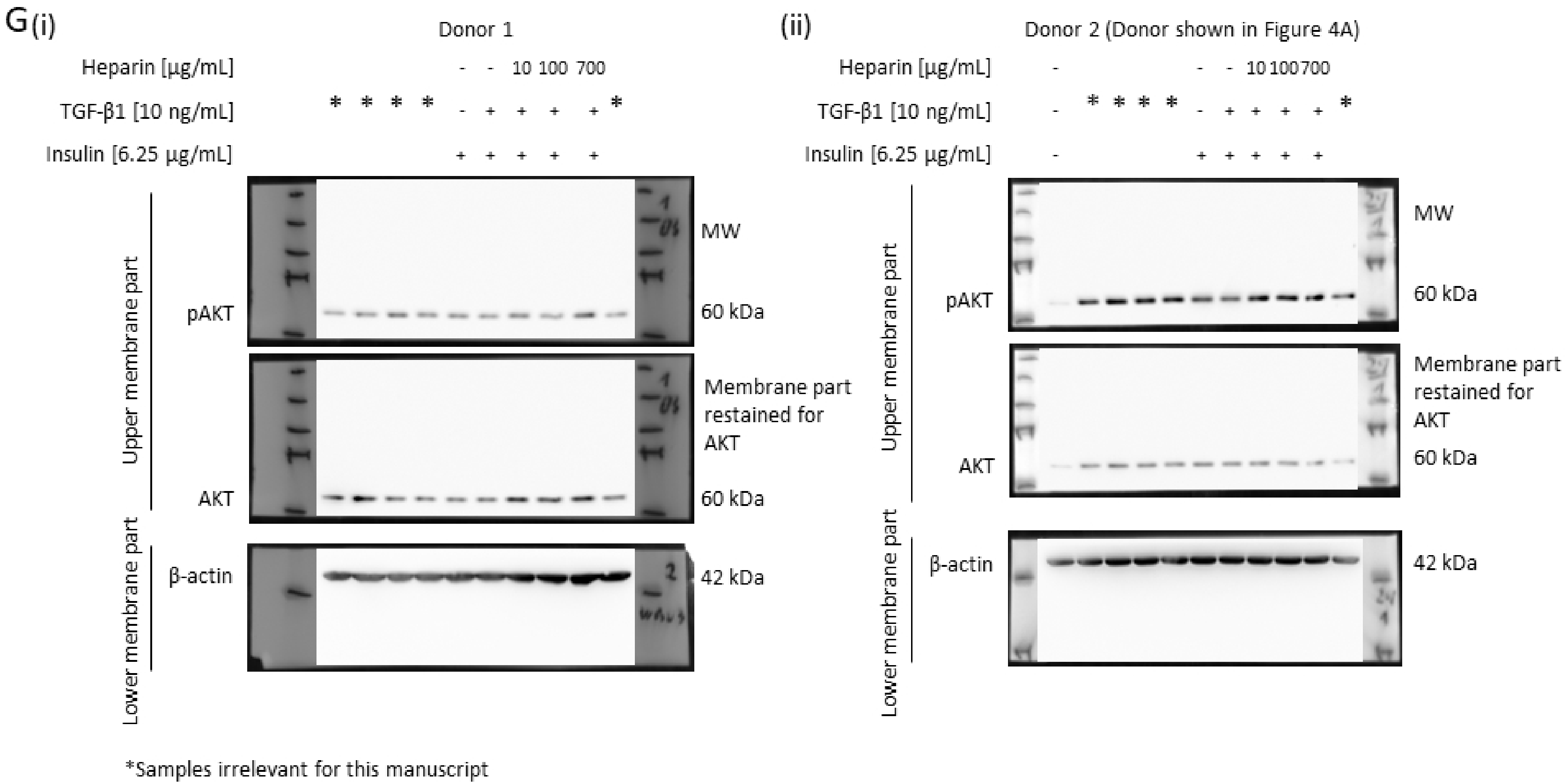


**Supplementary Material S1.** Western blots included in this study shown as full uncropped images.

(iii)

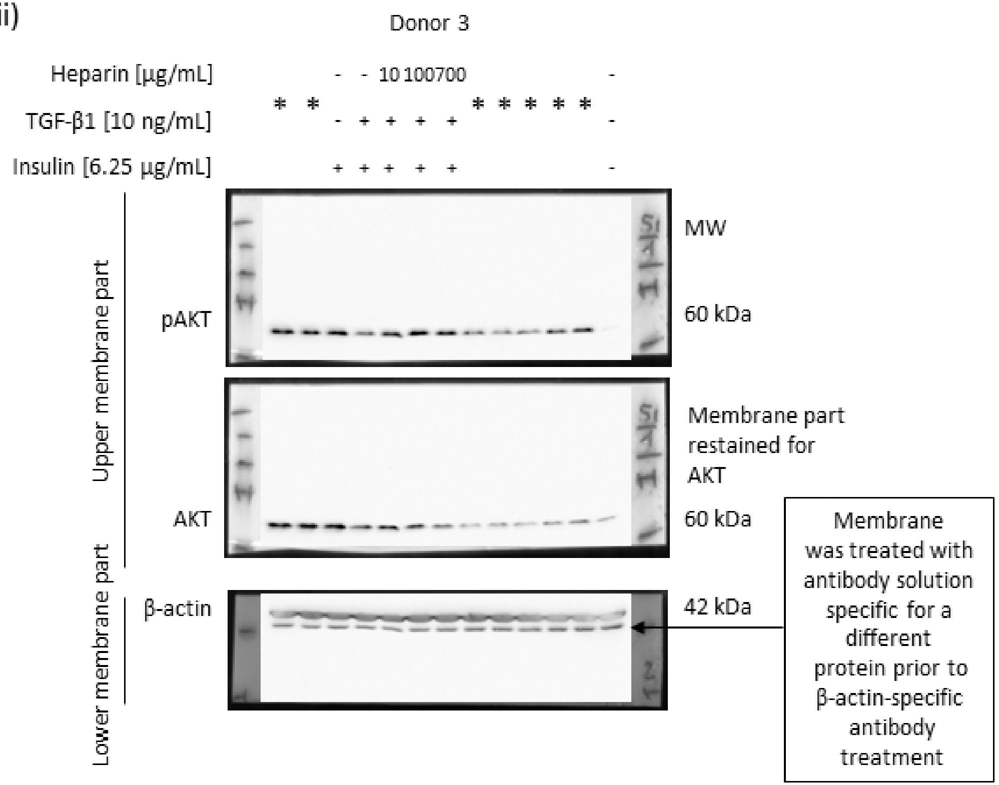


*Samples irrelevant for this manuscript

**Supplementary Material S1.** Western blots included in this study shown as full uncropped images.

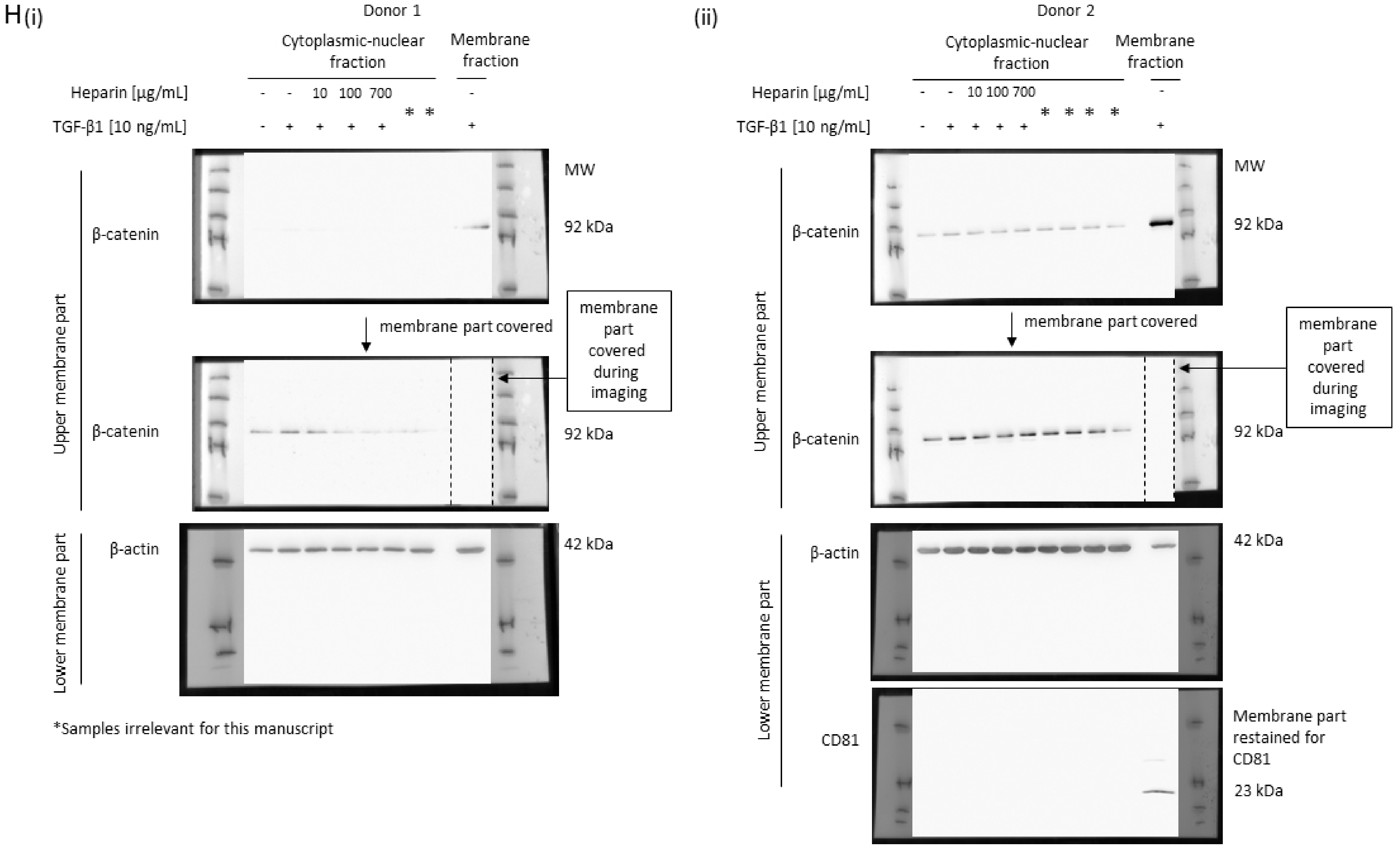


**Supplementary Material S1.** Western blots included in this study shown as full uncropped images.

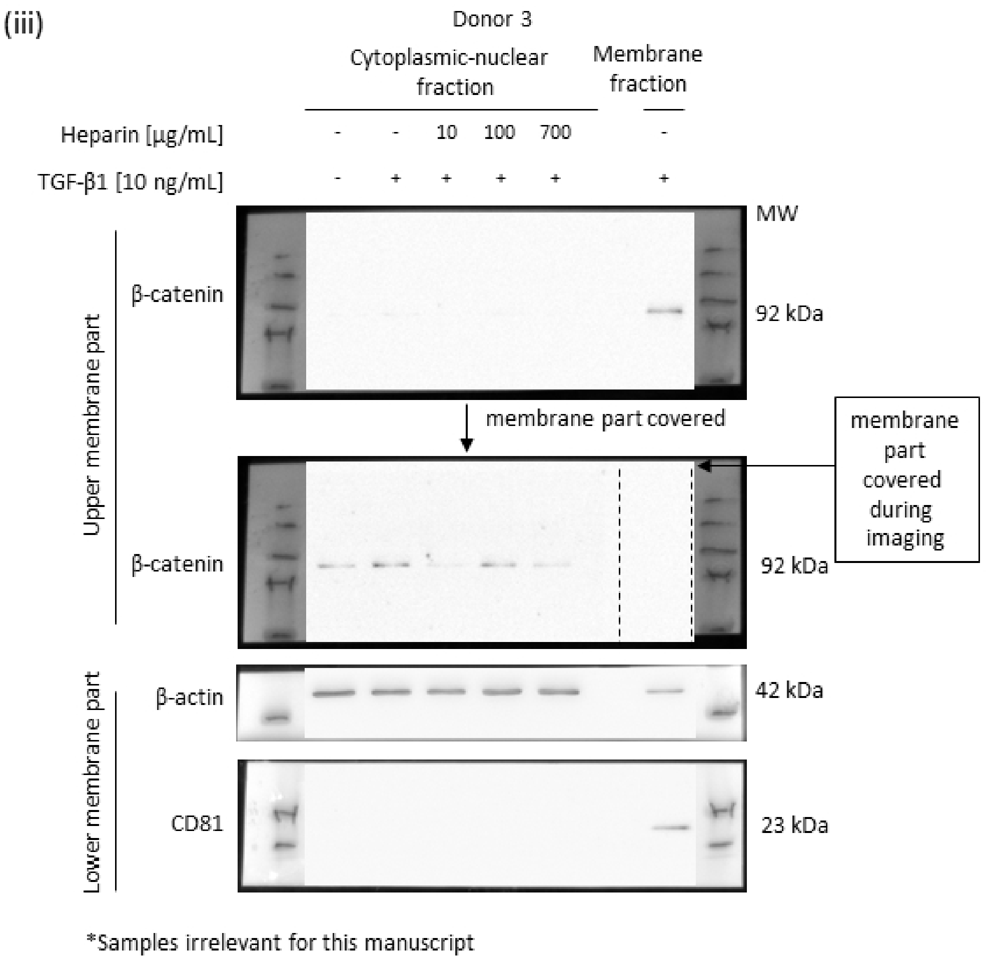


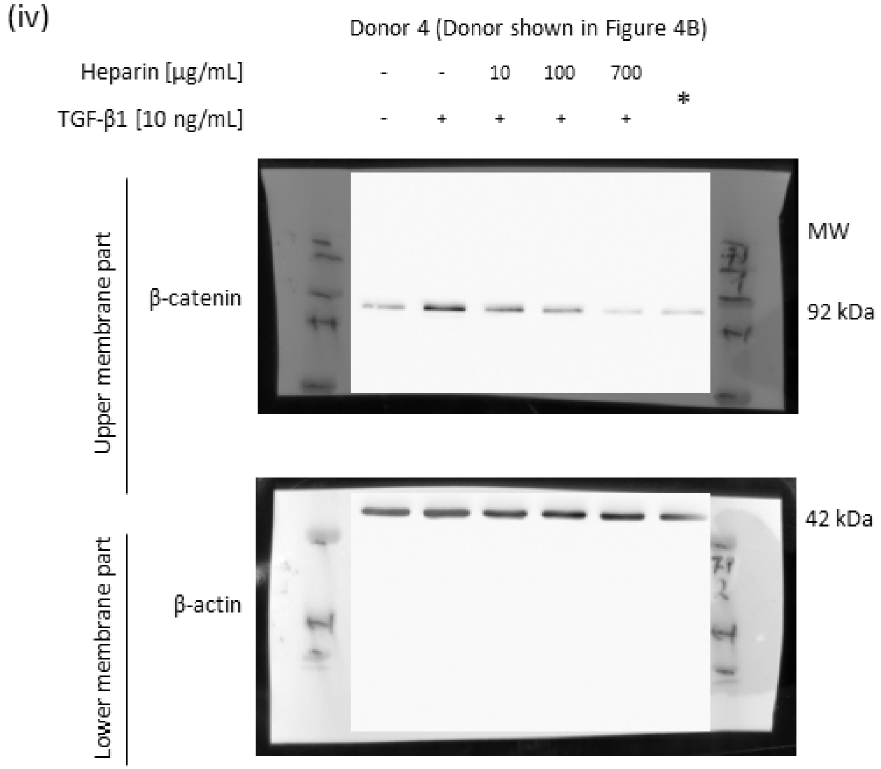


**Supplementary Material S1.** Western blots included in this study shown as full uncropped images.

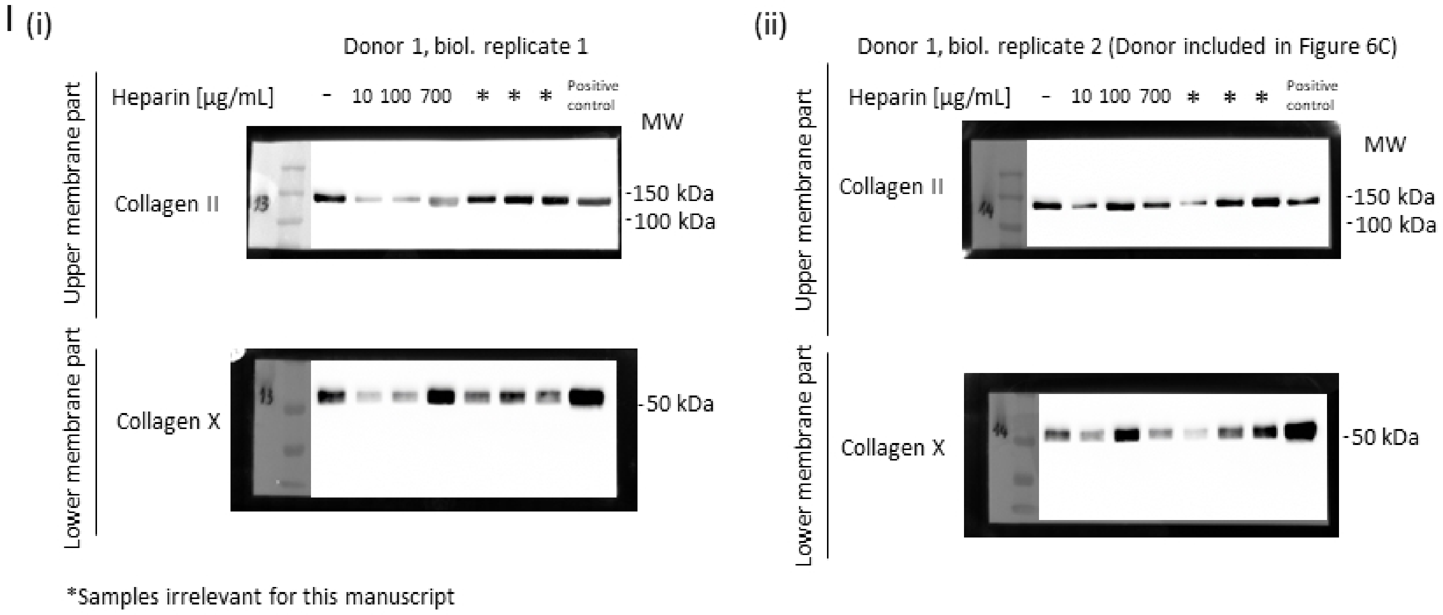


**Supplementary Material S1.** Western blots included in this study shown as full uncropped images.

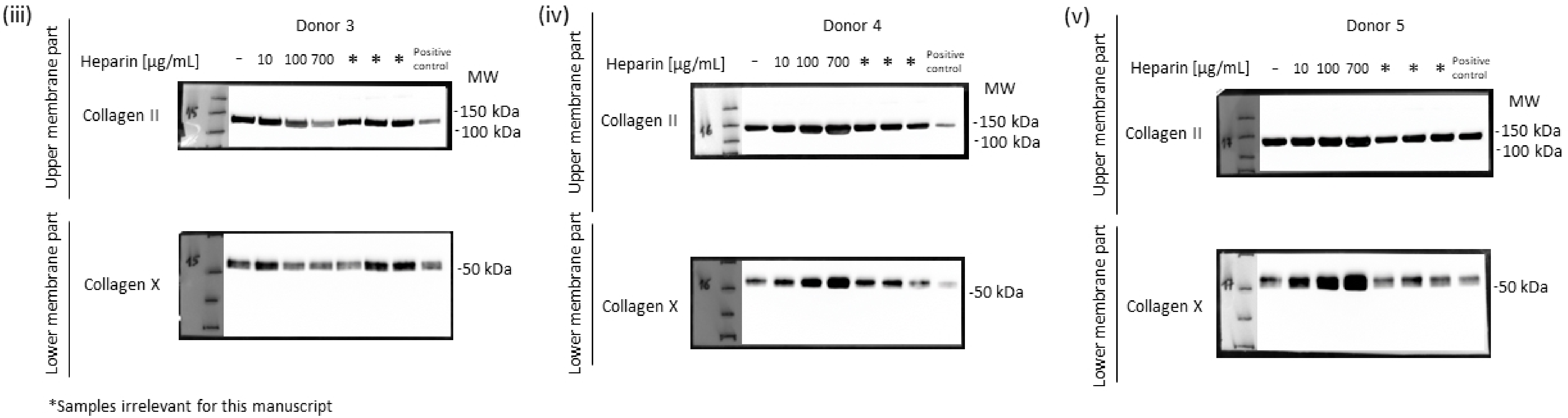


**Supplementary Material S1.** Western blots included in this study shown as full uncropped images.
