## Supplementary Table S1 for "Dichotomous SMAD2/3 regulation and selective anti-hypertrophic activity of heparin during in vitro chondrogenesis of mesenchymal stromal cells"

Supplementary Table S1: Alphabetical list of primer sequences utilized for qPCR analysis.

| Gen | Forward | Reverse |
| --- | --- | --- |
| ALPL | 5 ́-CACCAACGTGGCTAAGAATG-3 ́ | 5 ́-TCAGCTGGATGGCCACATC- 3 ́ |
| ACAN | 5 ́-GGAACCACTTGGGTCACG-3 ́ | 5 ́-GCACATGCCTTCTGCTT-3 ́ |
| COL10A1 | 5 ́-TTTACGCTGAACGATACCAAA-3 ́ | 5 ́-TTGCTCTCCTCTTACTGCTAT-3 ́ |
| COL2A1 | 5 ́-TGGCCTGAGACAGCATGAC-3 ́ | 5 ́-AGTGTTGGGAGCCAGATTGT-3 ́ |
| CPSF6 | 5 ́-AAGATTGCCTTCATGGAATTGAG-3 ́ | 5 ́-TCGTGATCTACTATGGTCCCTCTCT-3 ́ |
| GLI1 | 5 ́-TGCAGTAAAGCCTTCAGCAATG-3 ́ | 5 ́-TTTTCGCAGCGAGCTAGGAT-3 ́ |
| HPRT | 5 ́-AAGGGTGTTTATTCCTCATGGA-3 ́ | 5 ́-CCTCCCATCTCCTTCATCAC-3 ́ |
| IBSP | 5 ́-CAGGGCAGTAGTGACTCATCC-3 ́ | 5 ́-TCGATTCTTCATTGTTTTCTCCT-3 ́ |
| IHH | 5 ́-CGACCGCAATAAGTATGGAC-3 ́ | 5 ́-GGTGAGCGGGTGTGAGTG-3 ́ |
| MEF2C | 5 ́-GTATGGCAATCCCCGAAACT-3 ́ | 5 ́-ATCGTATTCTTGCTGCCTGG -3 ́ |
| PTH1R | 5 ́-GGTGAGGTGGTGGCTGT-3 ́ | 5 ́-AGCATGAAGGACAGGAAC-3 ́ |
| SOX9 | 5 ́-GTACCCGCACTTGCACAAC-3 ́ | 5 ́-TCGCTCTCGTTCAGAAGTCTC-3 ́ |
